# A strict requirement in proteasome substrates for spacing between ubiquitin tag and degradation initiation elements

**DOI:** 10.1101/2023.08.08.552540

**Authors:** Caroline Davis, B. L. Spaller, Erin Choi, Joseph Kurrasch, Haemin Chong, Suzanne Elsasser, Daniel Finley, Andreas Matouschek

**Author notes:** Correspondence and requests for materials should be addressed to A.M.

## Abstract

Proteins are typically targeted to the proteasome for degradation through the attachment of ubiquitin chains and the proteasome initiates degradation at a disordered region within the target protein. Yet some proteins with ubiquitin chains and disordered regions escape degradation. Here we investigate how the position of the ubiquitin chain on the target protein relative to the disordered region modulates degradation and show that the distance between the two determines whether a protein is degraded efficiently. This distance depends on the type of the degradation tag and is likely a result of the separation on the proteasome between the receptor that binds the tag and the site that engages the disordered region.

## Introduction

The Ubiquitin Proteasome System (UPS) is the major pathway of regulated protein degradation in eukaryotes. It clears damaged and misfolded proteins from cells, digests foreign proteins as part of the adaptive immune response, and controls the concentration of regulatory proteins involved in diverse cellular processes (Finley *et al*, 2016). The 26S proteasome is the proteolytic machine at the center of the UPS. Proteins are primarily targeted to the proteasome by the covalent attachment of one or more ubiquitin molecules to lysine residues within target proteins by the sequential action of E1, E2, and E3 enzymes. The process forms polyubiquitin chains in which the ubiquitin moieties are linked through isopeptide bonds between the C-terminus of one ubiquitin and one of the seven Lys residues or the N-terminal Met residue of another ubiquitin (Komander & Rape, 2012a; Li & Ye, 2008; Ohtake & Tsuchiya, 2017). Once a ubiquitinated protein is recognized by the proteasome, degradation initiates at a stretch of disordered amino acids within the substrate (Prakash *et al*, 2004b; Tomita & Matouschek, 2019b; Schmidt *et al*, 2005; Davis *et al*, 2021; Sahu & Glickman, 2021; Bard *et al*, 2018a; Elsasser & Finley, 2005; Whiteley *et al*, 2021, 2017; Saeki, 2017).

The proteasome is composed of a cylindrical 20S Core Particle (CP) and a 19S Regulatory Particle (RP) that caps either or both ends of the CP (Bard *et al*, 2018b). Access to the proteolytic chamber in the CP is controlled by the RP, which recognizes substrates and transfers them to the CP for degradation. Substrates travel from the RP to the CP via the substrate translocation channel, a passage that, because it is narrow, allows entry of unfolded polypeptides but excludes folded domains. This channel within the RP and is formed by a heterohexameric ring of ATPases of the AAA^+^ family, known as Rpt1-Rpt6. At least three subunits of the RP recognize ubiquitin: Rpn10, Rpn13, and Rpn1 (Deveraux *et al*, 1994; Husnjak *et al*, 2008b; Schreiner *et al*, 2008b; Shi *et al*, 2016a). The degradation rates of ubiquitinated substrates typically show the strongest dependence on Rpn10, which is situated closest to the entrance of the substrate translocation channel. Rpn13 is located at the top of the RP, somewhat further from the entrance channel than Rpn10 (Husnjak *et al*, 2008b; Schreiner *et al*, 2008b; Sakata *et al*, 2012), while Rpn1 is located on the opposite side of the entrance channel relative to Rpn10.

Rpn10, Rpn13, and Rpn1 can also bind UBL domains of UBL-UBA proteins (Husnjak *et al*, 2008b; Shi *et al*, 2016a; Schreiner *et al*, 2008b; Elsasser *et al*, 2002b). UBL-UBA proteins contain an N-terminal UBL domain connected by a flexible linker to one or more copies of a Ubiquitin-Associating (UBA) domain, which binds ubiquitin. Hence, UBL-UBA proteins are thought to act as extrinsic substrate receptors by binding polyubiquitin through their UBA domain(s) and the proteasome through their UBL domain (Bertolaet *et al*, 2001; Wilkinson *et al*, 2001; Chen *et al*, 2001; Rao & Sastry, 2002; Raasi *et al*, 2004; Elsasser *et al*, 2004; Elsasser & Finley, 2005; Whiteley *et al*, 2021, 2017; Zientara-Rytter & Subramani, 2019; Saeki, 2017). Some UBL-UBA proteins are required for the degradation of select natural proteins, though the mechanism of action remains unclear (Verma *et al*, 2004).

Recognition by ubiquitin receptors is not sufficient for substrate degradation. The proteasome must also physically engage the target protein at a segment of unstructured amino acids to initiate degradation (Prakash *et al*, 2004b; Takeuchi *et al*, 2006b). This initiation region inserts into the substrate translocation channel and is engaged by hydrophobic pore loops (pore-1 loops) of the ATPase ring. The initiation region must be of the appropriate length and composition for effective degradation (Tomita & Matouschek, 2019b; Yu *et al*, 2016a; Fishbain *et al*, 2015b; Inobe *et al*, 2011b; Yu *et al*, 2016d; Fishbain *et al*, 2011b). Initiation regions at the C-terminus of a protein must be some 30 residues in length for efficient degradation, but internally located initiation regions must be much longer (Prakash *et al*, 2004a; Inobe *et al*, 2011a; Takeuchi *et al*, 2006a; Tomita & Matouschek, 2019a; Fishbain *et al*, 2011a).

The proteasome must be able to precisely distinguish between its substrates and other ubiquitinated proteins in the cell to avoid promiscuous degradation. Intriguingly, ubiquitination and the presence of an initiation region do not always lead to rapid or efficient degradation (Martinez-Fonts *et al*, 2020b; Flick *et al*, 2004). How then does the proteasome select its substrates? Perhaps the arrangement of ubiquitin modification sites and the initiation region within some substrates prevents their simultaneous recognition by ubiquitin and initiation region receptors of the proteasome, preventing effective degradation (Inobe *et al*, 2011a). The initiation region binding site is situated within the substrate translocation channel, while the three ubiquitin receptors of the RP are located at distinct distances and directions away from the channel (Finley *et al*, 2016; Bard *et al*, 2018b). It stands to reason that an initiation region must at least be long enough to bridge the distance between the entrance to the translocation channel and the pore loops for effective engagement.

Here we determine how the position of the initiation region relative to a ubiquitin tag affects substrate selection by the proteasome by measuring the degradation of defined model substrates in which an initiation region is placed varying distances from the ubiquitin chain on the substrate. Our data suggest that the receptor that mediates substrate binding determines whether the proteasome can initiate degradation, presumably due to its distance from the pore loops within the substrate translocation channel. These observations provide new insight into the mechanism of substrate selection by the proteasome.

## Results

### Ubiquitin chain length and initiation

We first sought to determine how initiation region length affects the degradation of substrates possessing the canonical degradation-targeting signal (Ub_4_K48) (Thrower *et al*, 2000). To this end, we built substrates based on a central fluorescent protein domain (derived from GFP) with an N-terminal in-frame ubiquitin domain and a C-terminal initiation region (Fig 1a). We attached defined K48-linked ubiquitin chains to the N-terminal ubiquitin domain in purified base protein and measured fluorescence over time in the presence of purified proteasome, as described previously (see methods) (Martinez-Fonts & Matouschek, 2016; Martinez-Fonts *et al*, 2020b). Here we refer to the initiation region as a tail because of its location at the C-terminus of the fluorescent model protein. Tails were either 18 or 38 amino acids in length (Appendix Table S3). RP and CP were affinity purified from *Saccharomyces cerevisiae* with high salt washes to remove co-purifying proteasome-interacting proteins (Shi *et al*, 2016a; Leggett *et al*, 2005). The proteasome was reconstituted by mixing CP and RP in a 1:2 molar ratio.

**Figure 1.**
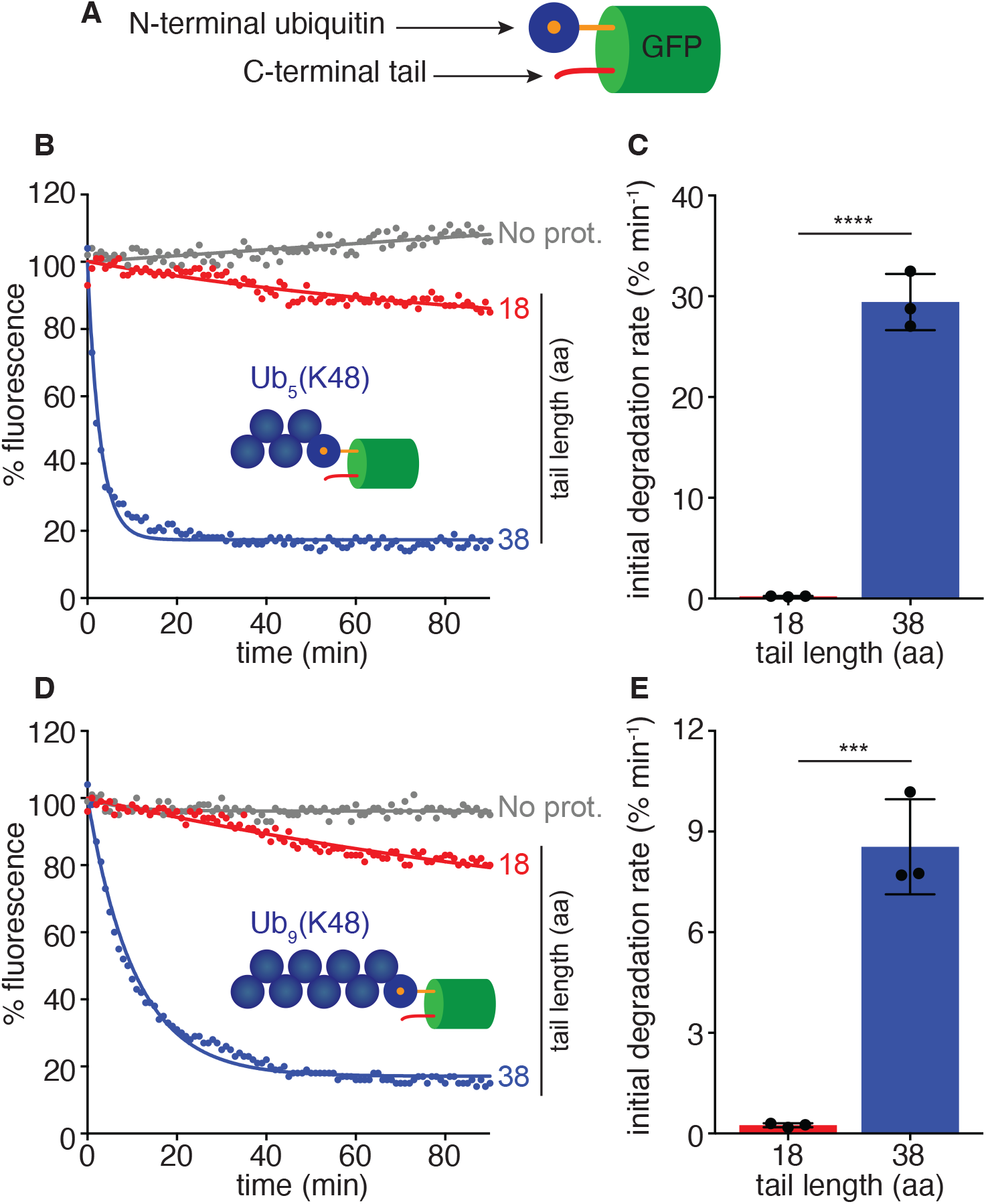
Ubiquitin chain length cannot compensate for initiation region length. A Schematic of *in vitro* ubiquitinated model substrates. Substrates consist of an N-terminal in-frame ubiquitin domain (dark blue with orange center) that serves as the attachment point for ubiquitin chains of precise length, followed by a GFP domain (green), and a C-terminal initiation region referred to as a tail (red). B *In vitro* degradation kinetics of Ub_5_(K48)-tagged GFP substrates with 18 (red) or 38 (blue) amino acid (aa) length tails under single turnover conditions (5 nM substrate, 25 nM wildtype proteasome) plotting substrate fluorescence as a percentage of the initial fluorescence as a function of time in minutes. No proteasome (no prot) control included. C Initial degradation rates of substrates with the indicated tail from B calculated by fitting kinetics to single exponential decay. D *In vitro* degradation kinetics of Ub_9_(K48)-tagged GFP substrates with 18 (red) or 38 (blue) amino acid (aa) length tails under single turnover conditions (5 nM substrate, 25 nM wildtype proteasome) plotting substrate fluorescence as a percentage of the initial fluorescence as a function of time in minutes. E Initial degradation rates of substrates with the indicated tail from D calculated by fitting kinetics to single exponential decay. Data information: In (B,D), experiments were performed in triplicate, one representative kinetic experiment shown. In (C, E) data represent the mean from three experiments and error bars represent the standard deviations (SD). **** P < 0.0001 and *** P < 0.001 (two-tailed unpaired t test). Proteasome types are described in Appendix Table S1.

In presence of excess proteasome, thus under single turnover conditions, substrate with the longer tail was degraded rapidly, whereas that with a short tail remained stable (Fig 1b,c), as observed previously (Inobe *et al*, 2011b; Verhoef *et al*, 2009; Prakash *et al*, 2004b; Takeuchi *et al*, 2006b). Assuming that the initiation region is too short to reach the pore loops in the substrate translocation channel, it is possible that increasing the length of the ubiquitin chain would change how the substrate binds Ub/UBL receptors, perhaps positioning the short tail for productive engagement by the proteasome. However, increasing the length of the ubiquitin chain from five to nine ubiquitin moieties did not enhance degradation of the short-tail substrate (Fig 1d,e). Interestingly, the longer ubiquitin chain reduced degradation of the longer-tail substrate, a pattern observed in previous steady-state experiments (Martinez-Fonts *et al*, 2020a). Thus, extending the ubiquitin chain does not compensate for a short initiation region.

### Moving the initiation region away from the ubiquitin chain

We then asked whether moving the tail further from the ubiquitin chain by altering the base substrate itself would increase degradation. We inserted stable α-helices of varying length (Yu *et al*, 2016a; Sivaramakrishnan *et al*, 2008) between GFP and the tail to move the tail further from the ubiquitin chain (Fig 2a). These α-helices, referred to here as linkers, were derived from mouse myosin VI and were found to be stable even in isolation in the absence of stabilizing tertiary interactions (Yu *et al*, 2016a; Sivaramakrishnan *et al*, 2008) Degradation of the substrates increased sharply with the addition of a linker of only 9 residues. Extension of the linker to 19 residues accelerated degradation further to its peak rate, after which lengthening the linker further to 30 residues or more suppressed degradation again. The linkers were not functioning as initiation regions since removing the tail inhibited degradation substantially (Fig EV1a), presumably because the amino acid sequences of the linkers are charged and compositionally biased (Table S4), which prevents proteasomal recognition (Yu *et al*, 2016a). These results suggest that efficient degradation of a ubiquitinated substrate requires placing the initiation region at the right distance from the ubiquitin chain; as placing the ubiquitin chain too close or too far relative the initiation region strongly suppresses degradation.

**Figure 2.**
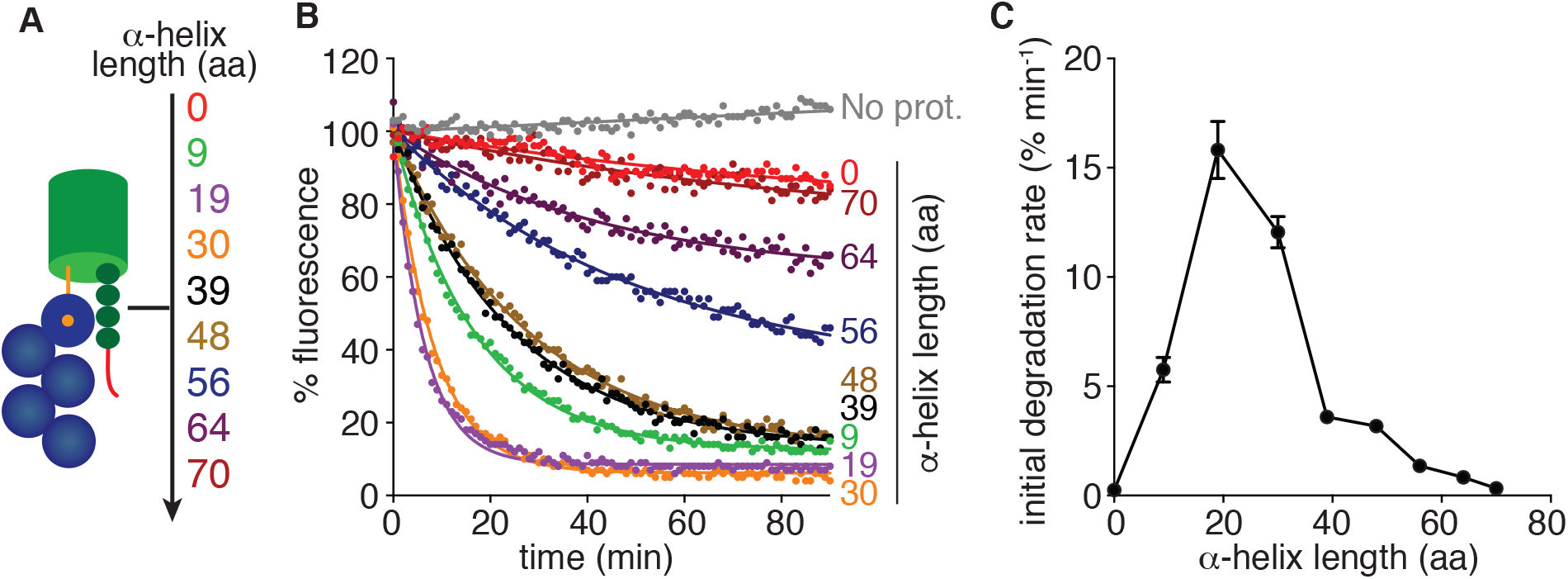
The initiation region must be placed at a distance from a ubiquitin chain to trigger efficient degradation. A Schematic of *in vitro* ubiquitinated model spacing substrates. Substrates contain an N-terminal Ub_5_(K48) tag (dark blue circles), a C-terminal 15 tail (red), and varying amino acid (aa) lengths of α-helices inserted between the GFP and tail (dark green circles). The amino acid lengths of the α-helices used are shown as numbers. B *In vitro* degradation kinetics of spacing substrates from A under single turnover conditions (5 nM substrate, 25 nM wildtype (WT) proteasome) plotting substrate fluorescence as a percentage of the initial fluorescence as a function of time in minutes. The amino acid (aa) length of the α-helix is shown to the right of the corresponding curve. C Initial degradation rates of substrates from B calculated by fitting kinetics to single exponential decay or linear equation. Graph plots the rate as a function of amino acid length of the α-helix. Data information: In (B), experiments were performed in triplicate, one representative kinetic experiment shown. In (C) data are plotted as a mean calculated from at least three experiments +/- the SD.

### Degradation of model substrates mediated by specific proteasomal ubiquitin receptors

The three ubiquitin receptors of the RP are located at different distances and directions from the entrance to the substrate translocation channel, which houses the pore-1 loops responsible for engaging initiation regions (see discussion) (Lander *et al*, 2012; Lasker *et al*, 2012; Beck *et al*, 2012; Unverdorben *et al*, 2014; Peña *et al*, 2018). Rpn10 is adjacent and closest to the translocation channel (65-95 Å from pore-1 loops), Rpn1 is located on the opposite side of the RP from Rpn10 (∼100 Å from pore-1 loops), but at its outer surface furthest from the translocation channel, and Rpn13 is at the top of the RP at a somewhat shorter distance from the translocation channel than Rpn1 (∼107 Å from pore-1 loops). Thus, the optimal linker length between ubiquitination site and initiation region could be characteristic of the particular ubiquitin receptor, on the assumption that concurrent binding of the ubiquitin chain by the receptor and of the initiation region by the pore-1 loops is required for degradation. To test this model, we engineered proteasomes in which only a single identified receptor was functional, thereby limiting substrate binding to one receptor. Each receptor was mutated so as to specifically abolish ubiquitin binding, using well-characterized substitution mutants: Rpn10 in its UIM element (*rpn10-UIM)* (Elsasser *et al*, 2004); Rpn13 in its PRU domain (*rpn13-PRU*) (Schreiner *et al*, 2008b; Husnjak *et al*, 2008b); and Rpn1 in its T1 and T2 sites (*rpn1-ARR-AKAA*) (Shi *et al*, 2016a). Here we refer to proteasomes by the single receptor that retains Ub/UBL binding (Appendix Table S1). Proteasomes in which ubiquitin binding to all three receptors is attenuated are called Quadruple Mutant (QM).

Rpn10 and wildtype proteasomes degraded ubiquitinated substrates equally well (Compare Fig 3a and Fig 2b) suggesting that K48-linked ubiquitin chains targeted the substrates to degradation principally through Rpn10, consistent with previous findings (Martinez-Fonts *et al*, 2020b). Degradation was strongly reduced on Rpn13 proteasome (Fig 3b), Rpn1 proteasome (Fig 3c), and QM proteasome (Fig 3d), though not completely abolished. The narrow width of the peak in degradation rates (Fig 3a) relative to linker length seen for degradation mediated by Rpn10 suggests that the spacing requirements are surprisingly well defined, considering that the UIM is thought to be flexibly connected to the proteasome. In the absence of Rpn10 Ub/UBL binding, the residual degradation appears less dependent on the linker length and spacing, perhaps because the residual degradation is mediated by multiple receptors with differing linker length optimal (Paraskevopoulos *et al*, 2014; Lam *et al*, 2002; Boughton *et al*, 2021).

**Figure 3.**
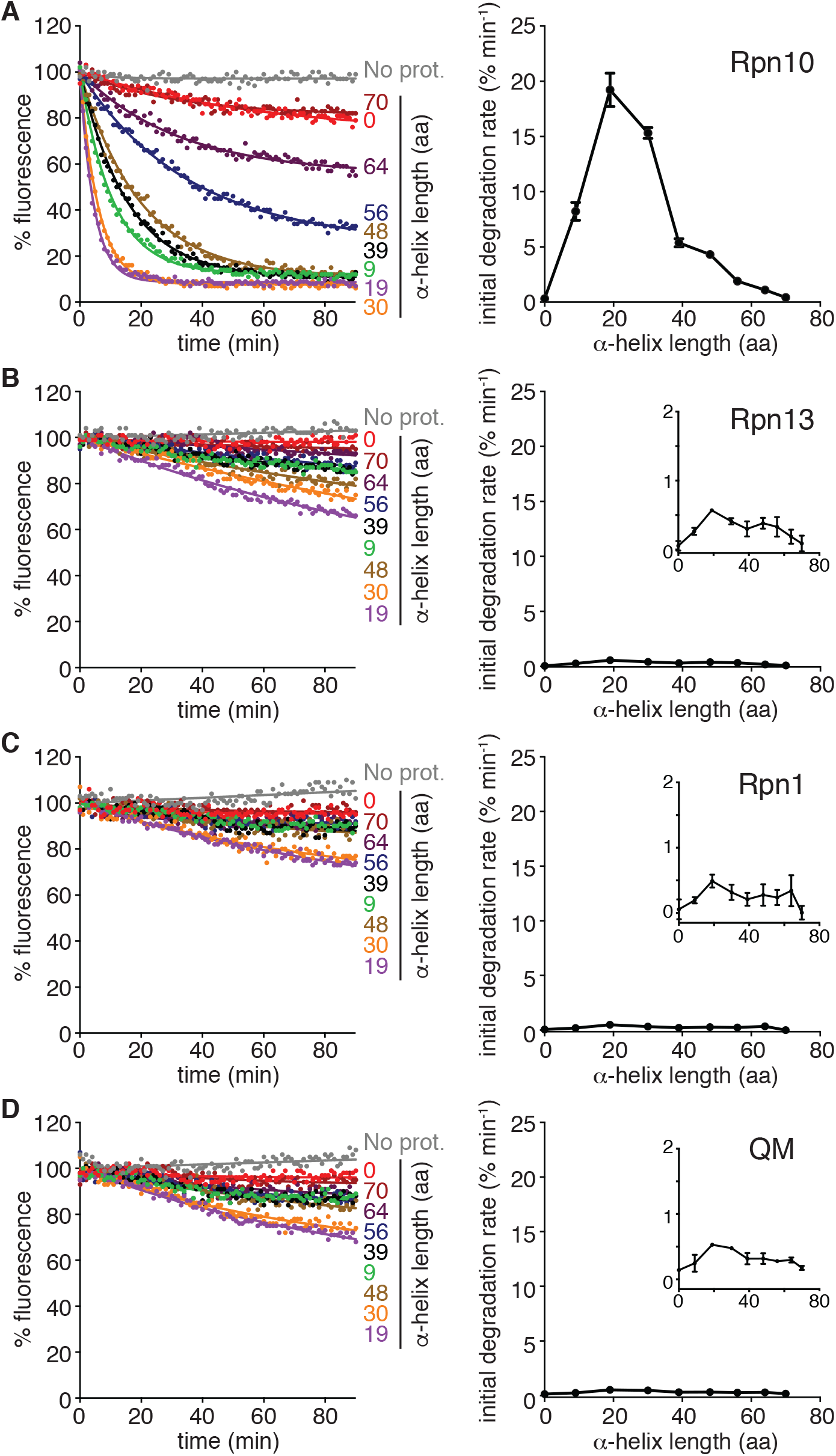
Optimal spacer lengths of proteasomal ubiquitin receptors. A-D (Left) *In vitro* degradation kinetics of ubiquitinated model spacing substrates under single turnover conditions (5 nM substrate, 25 nM proteasome) by the indicated proteasome receptor mutant (Rpn10, Rpn13, Rpn1, QM). Graphs plot substrate fluorescence as a percentage of the initial fluorescence as a function of time in minutes. The amino acid (aa) length of the α-helix is shown to the right of the corresponding curve. (Right) Initial degradation rates of ubiquitinated spacing substrates from the left calculated by fitting kinetics to single exponential decay or linear equation. Graphs plot the rate as a function of amino acid length of the α-helix. Data information: In (A-D, left), experiments were performed in triplicate, one representative kinetic experiment shown. In (A-D), rate data are plotted as a mean calculated from at least three experiments +/- the SD. Proteasome types are described in Appendix Table S1.

### Degradation of model substrates with an UBL tag

UBL-UBA proteins are thought to function as extrinsic substrate receptors, and there are three in yeast (Rad23, Dsk2, Ddi1) (Bertolaet *et al*, 2001; Wilkinson *et al*, 2001; Chen *et al*, 2001; Rao & Sastry, 2002; Raasi *et al*, 2004; Elsasser *et al*, 2004; Husnjak *et al*, 2008a; Shi *et al*, 2016b; Schreiner *et al*, 2008a; Elsasser *et al*, 2002a). They can bind all three ubiquitin receptors of the proteasome through their UBL domain and bind ubiquitin chains through their UBA domains. To investigate degradation of substrates targeted to the proteasome via UBL-UBA proteins while bypassing the complexities of protein complex formation, we fused the UBL domain from yeast Rad23 directly to the N-terminus of GFP, and appended tails of 18 or 96 amino acids in length to the C-terminus (Fig 4a) (Appendix Table S3). As observed for substrate with ubiquitin chains, UBL substrate with the short tail resisted degradation whereas substrate with the longer tail was degraded readily (Fig 4b,c). We again moved the short tail away from the UBL tag by inserting linkers of increasing length after the GFP domain. At first, degradation accelerated slowly with linker length, but then increased sharply, peaking with a 56 residue linker, before decreasing again (Fig 4e,f). The linkers themselves did not allow the proteasome to engage the substrates as shown by control experiments in which removing the tail inhibited degradation (Fig EV1b).

**Figure 4.**
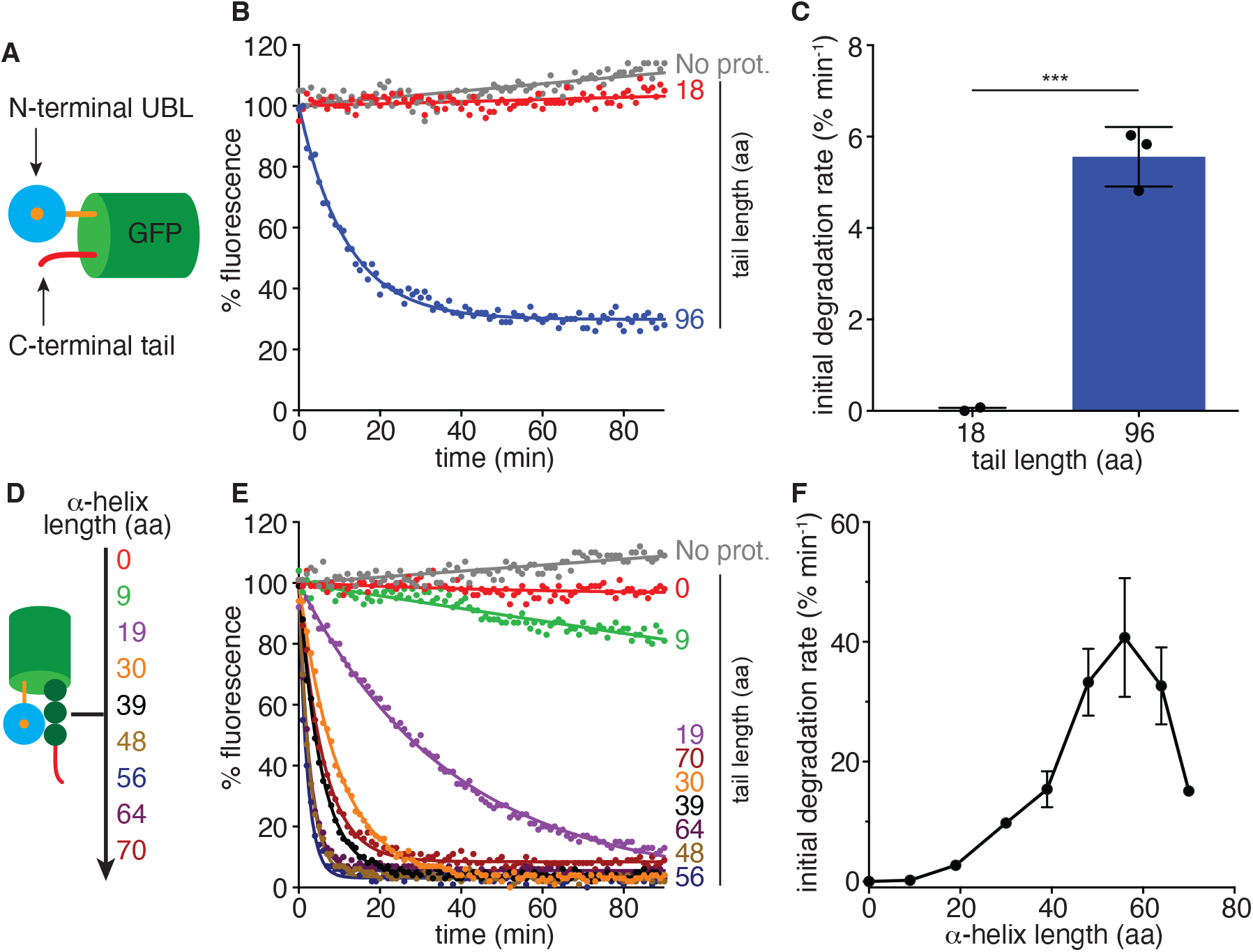
Optimal spacer length of UBL-tagged substrates is much longer than that of ubiquitin-tagged substrates. A Schematic of *in vitro* UBL model substrates. Substrates consist of the UBL domain of *S. cerevisiae* Rad23 (cyan with orange center) at the N-terminus, followed by a GFP domain (green), and a tail (red) at the C-terminus. B *In vitro* degradation kinetics of UBL-tagged GFP substrates with 18 (red) or 96 (blue) amino acid (aa) length tails under single turnover conditions (5 nM substrate, 25 nM wildtype (WT) proteasome) plotting substrate fluorescence as a percentage of the initial fluorescence as a function of time in minutes. C Initial degradation rates of substrates with the indicated tail from B calculated by fitting kinetics to single exponential decay. D Schematic of *in vitro* UBL model spacing substrates. Substrates contain an N-terminal UBL tag (cyan circle), a C-terminal 18 tail (red), and varying amino acid (aa) lengths of α-helices inserted between the GFP and tail (dark green circles). The amino acid lengths of the α-helices used are shown as numbers. E *In vitro* degradation kinetics of spacing substrates from E under single turnover conditions (5 nM substrate, 25 nM WT proteasome) plotting substrate fluorescence as a percentage of the initial fluorescence as a function of time in minutes. The amino acid length of the α-helix is shown to the right of the corresponding curve. F Initial degradation rates of substrates from E calculated by fitting kinetics to single exponential decay. Graph plots the rate as a function of amino acid length of the α-helix. Data information: In (B,E), experiments were performed in triplicate, representative kinetic experiment shown. In (C), data represent the means calculated from three experiments. Error bars represent the standard deviations (SD). *** P < 0.001 (two-tailed unpaired t test). In (F), data are plotted as a mean calculated from at least three experiments +/- the SD.

Degradation of UBL substrates required greater distance between the two parts of the degradation signal than the ubiquitin substrates. It also proceeded largely through different Ub/UBL receptors. Substrates with the UBL domain were degraded equally well on Rpn13 and wildtype proteasome (compare Fig 4e,f and Fig 5b), as expected (Martinez-Fonts *et al*, 2020b), and degradation was strongly reduced on Rpn10 (Fig 5a) and QM (Fig 5d) proteasomes. The slow degradation by Rpn10 proteasome (Fig 5a) appeared to be mostly through residual binding to the mutated Rpn13, because deleting the Rpn13 subunit entirely in the Rpn10 proteasome (i.e., Rpn10 (*Rpn13*Δ)) reduced degradation further (Fig EV2). The remaining degradation by Rpn10 (*Rpn13*Δ) proteasome was likely due to the UBL domain binding the proteasome elsewhere because replacing the UBL domain with a protein of similar size but unrelated in sequence (Barstar) stabilized the substrate completely, even when all Ub/UBL binding sites were functional in wildtype proteasome (Fig EV3). In comparison, Rpn1 proteasome degraded the UBL substrates almost as well as Rpn13 proteasome (Fig 5c), suggesting that Rpn1 can also serve as a UBL receptor for degradation. Degradation through Rpn13 and Rpn1 changed with separation between UBL domain and tail in similar ways, likely because the two receptors are located at similar distances from the pore-1 loops.

**Figure 5.**
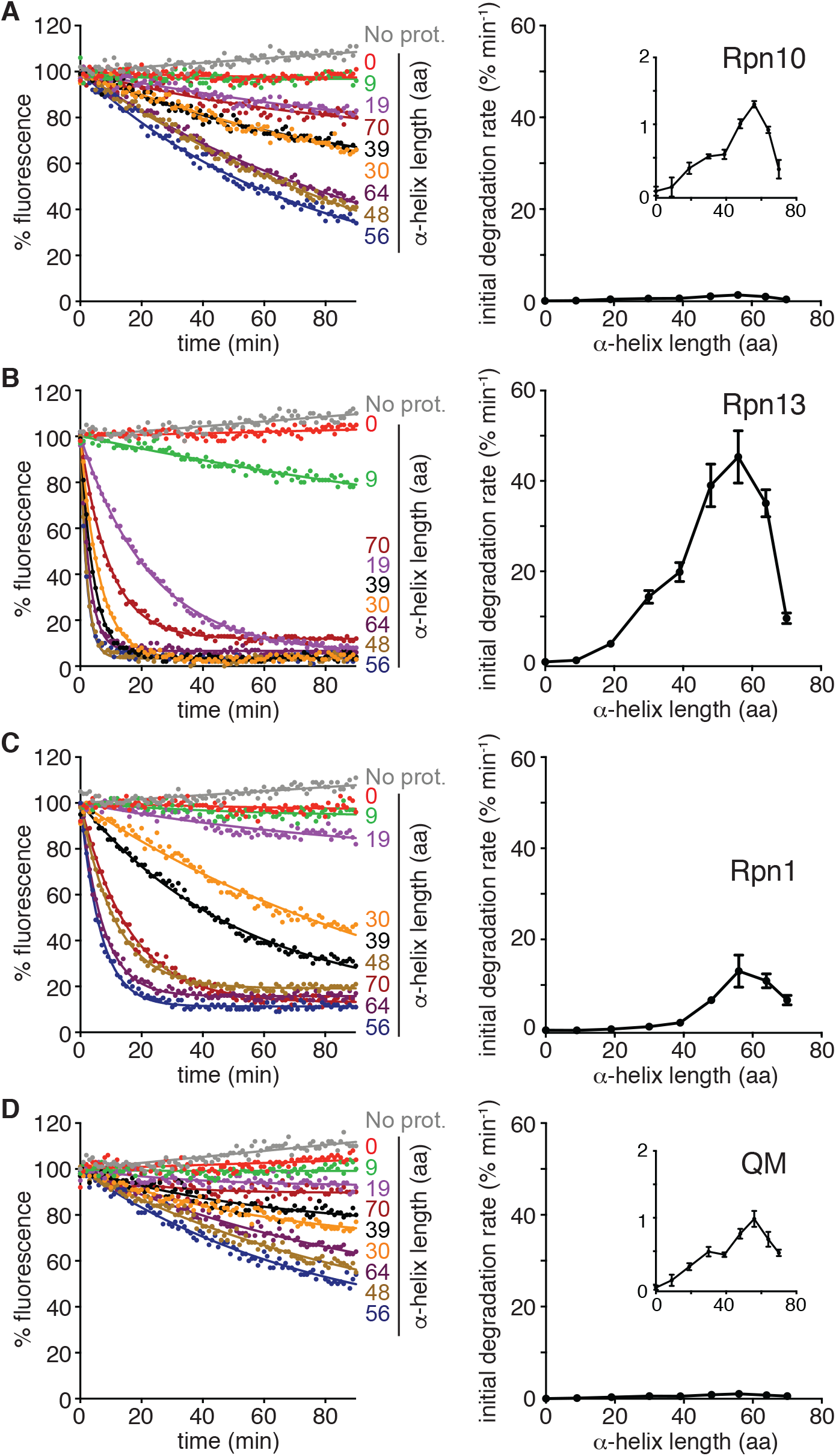
Degradation of UBL-tagged spacing substrates by proteasome receptor mutants reveals a preference for Rpn13. A-D (Left) *In vitro* degradation kinetics of UBL spacing substrates under single turnover conditions (5 nM substrate, 25 nM proteasome) by the indicated proteasome receptor mutant (Rpn10, Rpn13, Rpn1, QM). Graphs plot substrate fluorescence as a percentage of the initial fluorescence as a function of time in minutes. The amino acid (aa) length of the α-helix is shown to the right of the corresponding curve. (Right) Initial degradation rates of UBL spacing substrates calculated by fitting degradation kinetics to single exponential decay or linear equation. Graphs plot the rate as a function of amino acid length of the α-helix. Data information: In (A-D, left), experiments were performed in triplicate, one representative kinetic experiment shown. In (A-D, right), rate data are plotted as a mean calculated from at least three experiments +/- the SD. Proteasome types are described in Appendix Table S1.

### Degradation of model substrates in yeast

We asked next whether the distance between the proteasome-binding tags and the initiation region also controlled degradation in cells. We expressed model substrates analogous to those used in the *in vitro* experiments, alongside a stable red fluorescent reference protein and followed degradation by monitoring the fluorescence signals originating from the substrate and reference protein in single yeast cells by flow cytometry. We measured degradation rates by stopping protein synthesis with cycloheximide and monitored steady state accumulation by calculating the ratio of substrate fluorescence to reference protein fluorescence, as done previously (Yu *et al*, 2016b, 2016c). The yeast model substrates contained YFP instead of GFP and the UBL substrates were otherwise identical (Fig 6c). The ubiquitinated substrates differed further as ubiquitination was induced by an N-end degron attached to the N-terminus (Chau *et al*, 1989; Tomita *et al*, 2020) and so ubiquitin chains were attached on Lys residues in a 14-residue N-terminal tail attached to the YFP domain (Fig 6a). Nevertheless, degradation proceeded from the C-terminal tail as replacing a tail that is well recognized by the proteasome (Su9) with a tail that is poorly recognized (SP25) inhibited degradation (Fig. EV4a) (Yu *et al*, 2016b). Degradation of the ubiquitinated substrate depended on UBR1, which encodes the N-end rule E3, and neither ubiquitin nor UBL substrate relied on the autophagy machinery (Fig EV4b).

**Figure 6.**
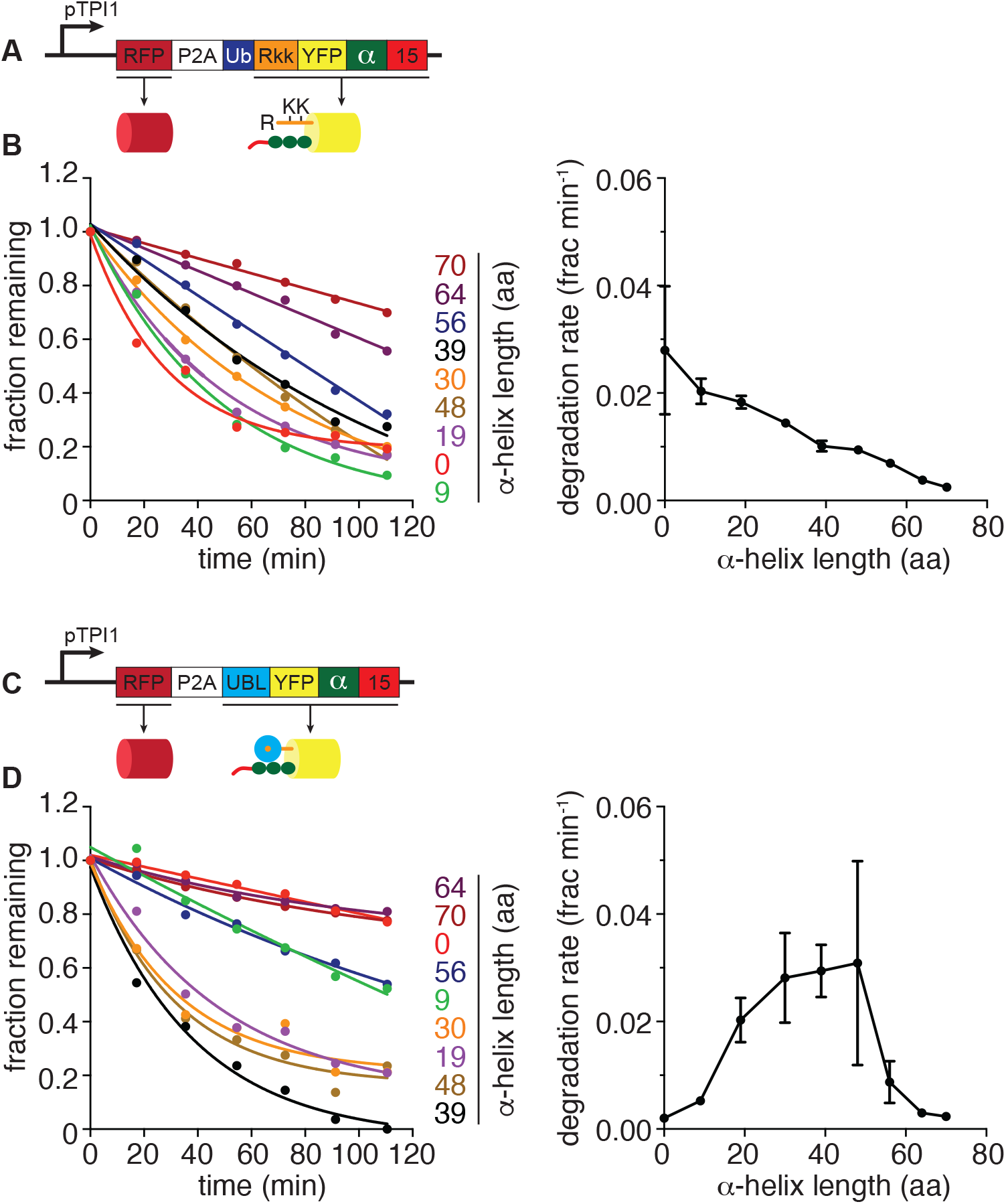
Observation of spacing differences in vivo. A,C Substrate design of fluorescence-based degradation assay in *S. cerevisiae*. DsRedExpress2 (RFP), followed by a ribosome skipping sequence (P2A), and model YFP substrate are expressed from a single promoter (pTPI1) to ensure equal production of the RFP reference protein and the YFP model substrates. (A) Ubiquitinated model substrates consist of an N-terminal in-frame ubiquitin domain (dark blue), followed by a modified N-end rule degron Rkk (orange), YFP (yellow), and a C-terminal tail (red). Upon expression in yeast, the ubiquitin domain is cleaved by ubiquitin hydrolases, exposing the modified N-end rule degron Rkk. Ubiquitin ligases recognize the degron and then ubiquitinate the lysine residues within the Rkk, which targets the substrate to the proteasome. (C) UBL model substrates consist of the UBL domain of *S. cerevisiae* Rad23 (cyan with orange center) at the N-terminus, followed YFP, and a C-terminal tail (red). B,D Yeast cells were treated with 125 µM cycloheximide to inhibit protein synthesis and YFP fluorescence was measured roughly every 20 minutes for approximately 120 minutes. B (Left) *In vivo* degradation kinetics of ubiquitinated YFP spacing substrates. Fluorescence plotted as a fraction of the initial fluorescence as a function of time and fitted to a single exponential decay or linear equation to calculate degradation rates. The amino acid (aa) length of the α-helix is shown to the right of the corresponding curve. (Right) Initial degradation rates of ubiquitinated YFP spacing substrates calculated by fitting degradation kinetics to single exponential decay or linear equation. Graph plots the rate as a function of amino acid length of the α-helix D (Left) *In vivo* degradation kinetics of UBL YFP spacing substrates. (Right) Initial degradation rates of UBL YFP spacing substrates calculated by fitting degradation kinetics to single exponential decay or linear equation. Graph plots the rate as a function of amino acid length of the α-helix Data information: In (B,D), experiments were performed in triplicate, one representative kinetic experiment shown. Degradation rate data are plotted as a mean calculated from at least three experiments +/- the SD Yeast strains are described in Appendix Table S5.

In yeast, degradation was most effective when the tail and ubiquitination site were placed next to each other, and degradation was faster for substrate without an additional linker inserted after the YFP domain, and gradually decreased as linker length increased (Fig 6b). *In vitro* degradation was most efficient with a linker of 19 residues inserted after the fluorescent protein (Fig 2c) but note that the N-end degron used in the yeast substrates inserts eleven unstructured residues before the fluorescent domain, which may function similarly to the short linkers used in the *in vitro* model substrates by positioning the tail a bit further from the ubiquitination site. The geometry of the ubiquitin chain in the two substrates also differs in the attachment point itself, where the chain is attached to a Lys residue in a folded domain *in vitro* and to Lys residue in an unstructured region in yeast.

Substrates targeted to the proteasome with a UBL domain were poorly degraded without a linker (Fig 6d) and degradation rates steadily improved as linker length increased until it plateaued, and then sharply decreased as linkers were lengthened past 56 residues. Substrates with very long linkers were degraded as poorly as substrate lacking any linker. Thus, degradation of UBL substrates in cells again followed the same pattern as *in vitro*. Degradation in yeast was somewhat more tolerant to separation and proceeded similarly well with linkers between 19 and 56 residues in length, roughly translated to 28.5 and 84 Å in length, respectively.

## Discussion

Controlling protein concentration through the precisely targeted degradation via the UPS is central to the regulation of many processes such as cell cycle progression, DNA repair, signaling cascades, and others. The proteasome must be able to degrade almost any protein but do so in a highly selective manner. Proteasomal degradation requires localization of the target protein to the proteasome, typically through a ubiquitin chain conjugated to the target protein, as well as a disordered region within the target protein of appropriate length and amino acid composition (Finley, 2009; Collins & Goldberg, 2017; Komander & Rape, 2012b; Tomita & Matouschek, 2019a). However, ubiquitination is also involved in regulating nonproteolytic processes (Groothuis *et al*, 2006), and intrinsically disordered regions are found throughout the entire proteome. Therefore, if ubiquitination and the existence of a disordered region were the only requirements for proteasomal degradation, most cellular proteins could be subject to proteasomal degradation at any given time. Instead, proteins can escape degradation, even with ubiquitin modifications and disordered regions, suggesting the degradation-targeting code is more nuanced than currently defined (Martinez-Fonts *et al*, 2020a; Inobe *et al*, 2011a). Here, we investigated a steric requirement for the degradation signal in proteasomal degradation.

We find that the position of the ubiquitin modification in a protein relative to the initiation region modulates its degradation *in vitro* and in yeast. The reason is likely found in the structure of the proteasome itself and the location of the ubiquitin binding receptors on the proteasome relative to the ATPase ring, where disordered initiation regions are inserted and physically engaged by the pore-1 loops. Previous biochemical studies show initiation regions must be of a certain minimal length to target for degradation (Inobe *et al*, 2011b; Yu *et al*, 2016d; Prakash *et al*, 2004b; Takeuchi *et al*, 2006b; Fishbain *et al*, 2015b; Lee *et al*, 2014; Yu *et al*, 2016a). For instance, C-terminal initiation regions must be approximately 30 residues in length for effective degradation (Prakash *et al*, 2004b; Inobe *et al*, 2011b; Takeuchi *et al*, 2006b; Tomita & Matouschek, 2019b). We find here that shorter initiation regions can trigger degradation if they are placed the required distance from a proteasome-binding tag. Thus, substrate degradation can be controlled by the spatial arrangement of the proteasome-binding tag and the initiation region on a protein.

Ancillary cellular factors such as UBL-UBA proteins and Cdc48 are believed to function in the UPS (Christianson & Ye, 2014; Elsasser *et al*, 2005; Saeki *et al*, 2002). UBL-UBA proteins are thought to shuttle substrates to the proteasome by binding ubiquitin chains and the proteasome through their UBA domain(s) and UBL domain, respectively (Rao & Sastry, 2002; Raasi *et al*, 2004; Elsasser *et al*, 2004; Saeki *et al*, 2002; Su & Lau, 2009; Elsasser & Finley, 2005; Zientara-Rytter & Subramani, 2019; Saeki, 2017; Whiteley *et al*, 2021, 2017). The UBL and UBA domain(s) are connected through long flexible linkers, which likely allow the domains, and any bound proteins, to explore many orientations relative to each other, contrasting the relative spatial rigidity of the Ub/UBL receptors and pore-1 loop positions on the proteasome (Fishbain *et al*, 2015a; Dantuma *et al*, 2009). Thus, UBL-UBA proteins could loosen the strict initiation region and ubiquitin chain spacing requirements observed *in vitro* by positioning of substrates so that disordered regions are close to the entrance to the substrate channel and the pore-1 loops. Cdc48, a homohexameric AAA+ ATPase, is thought to work upstream of the proteasome by unfolding proteins prior to delivery to the proteasome (Olszewski *et al*, 2019; Ji *et al*, 2022). Thus, in effect Cdc48 could lengthen the initiation region in substrates, allowing it to reach the pore-1 loops regardless of the placement of the ubiquitin chain in the folded state of the substrate protein.

Degradation of the fluorescent model substrates with different linkers followed similar patterns *in vitro* and in yeast (Figs 2, 4, and 6). The ubiquitin-dependent substrates both showed fastest degradation when the ubiquitin chain was close to the initiation region, whereas UBL substrates were degraded best when the initiation region was placed further from the UBL domain. The degradation profiles did differ in the details, most notably for the ubiquitinated substrates in which the constructs differed in the attachment sites of the ubiquitin chains. In addition, the peaks in degradation rates were noticeably broader in yeast than *in vitro*. Nevertheless, the fact that degradation of the substrates investigated here follows similar patterns *in vitro* and in yeast suggests that they interact directly with the proteasome, bypassing the accessory factors discussed above. We suspect that UBL-UBA proteins and Cdc48 did not recognize the model substrates because of some property of the ubiquitin chains such as their length (Bodnar *et al*, 2018), linkage, or branching pattern (Liu *et al*, 2017; French *et al*, 2021), or because they were not localized to the same subcellular location (Okeke *et al*, 2020). The mechanism by which such extrinsic cellular factors contribute to degradation of some natural proteins requires further investigation.

The Ub/UBL binding sites of Rpn13 and Rpn1 are approximately 107 Å and 100 Å away from the pore loops, which must engage the initiation region for degradation to proceed. The Ub/UBL binding site of Rpn10 has not been resolved, but the last resolved residue of the proteasome docked VWA domain of Rpn10 is 95 Å away from the pore-1 loops. The Ub/UBL binding domain is connected to the VWA domain by a flexible 20 amino acid long linker (Sakata *et al*, 2012), which may allow the Ub/UBL binding domain to approach the pore-1 loops and decrease the separation by some 30 Å. Substrates targeted to the proteasome by K48 linked ubiquitin chains are recognized primarily through Rpn10 and are degraded most rapidly when the ubiquitin chain is close to the initiation region. Substrates targeted to the proteasome by UBL domains are primarily recognized by Rpn13 and degraded most rapidly when the UBL domain is placed further from the initiation region compared to substrates with K48 linked ubiquitin chains. Thus, substrates targeted to the proteasome by ubiquitin chains and UBL domains require very different arrangements of proteasome-binding tag and initiation region for rapid degradation, probably because their Ub/UBL receptors on the proteasome are positioned at different distances form the pore-1 loops (Fig. 7).

**Figure 7.**
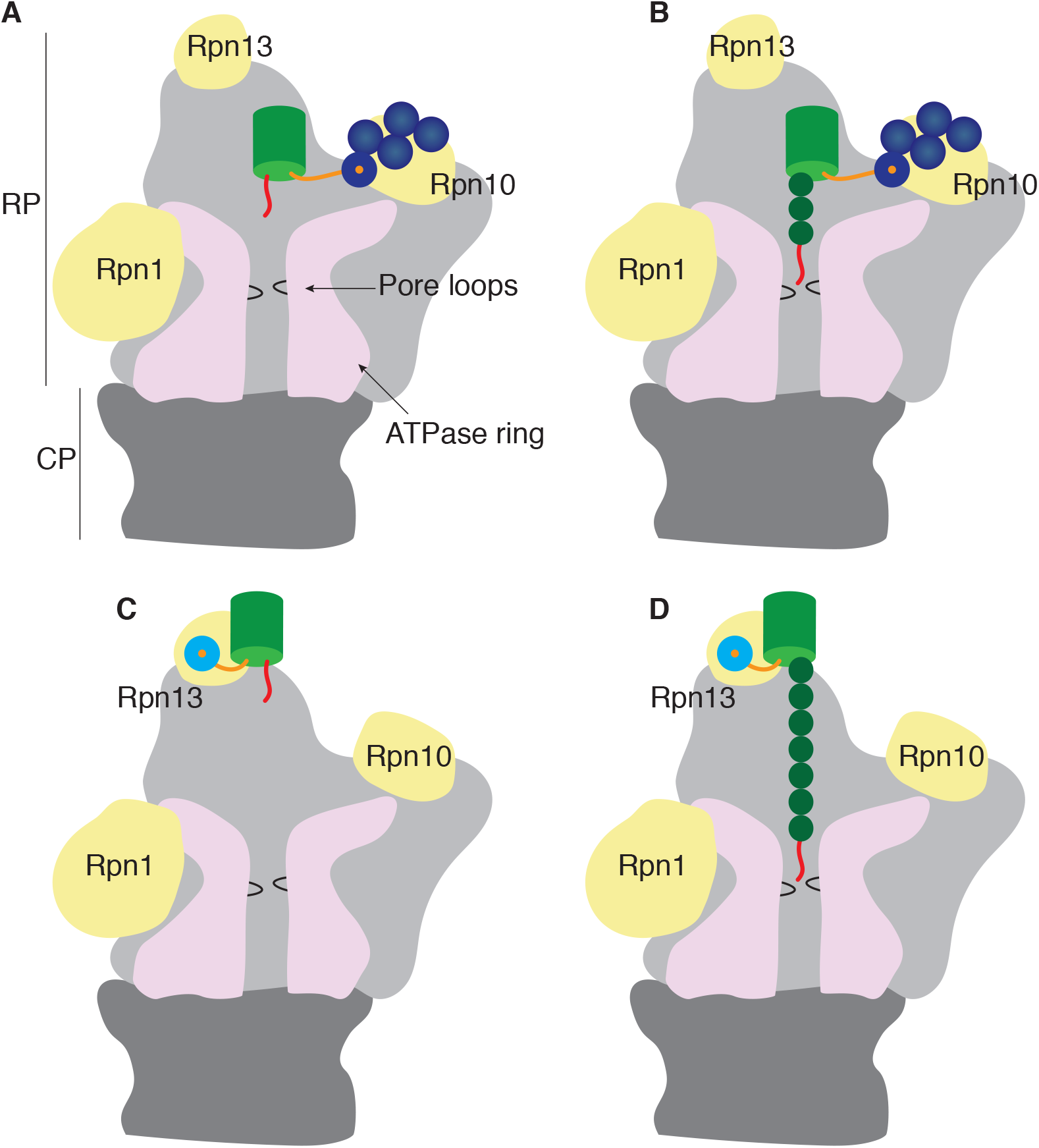
Model illustrating how position of initiation region relative to a proteasome-binding tag modulates engagement by the proteasome. A-D Cartoon of the proteasome highlighting the three known ubiquitin receptors (yellow) and the pore loops within the translocation channel formed by the ATPase ring (pink) of the RP. Substrates with variously arranged proteasome-binding tag and initiation region (red) superimposed on proteasome. Ubiquitin chains shown in dark blue and UBL domain shown in cyan. A The initiation region of a ubiquitinated substrate is unable to reach the pore loops when bound to Rpn10. B Degradation of substrate from A can be achieved when the initiation region is placed a short distance from the ubiquitin chain and by proxy Rpn10. C The initiation region of a substrate targeted to the proteasome through a UBL domain is unable to reach the pore loops when bound to Rpn13. D Degradation of substrate from C can be achieved when the initiation region is placed a far distance from the UBL domain and by proxy Rpn13.

The previously underappreciated importance of steric requirements in proteasome-mediated degradation has the potential to improve design of therapeutics that utilize the proteasome. Inducible degradation of endogenous cellular proteins has potential to address otherwise undruggable targets, offering new therapeutic applications (Donovan *et al*, 2020; Silva *et al*, 2019; Russ & Lampel, 2005). PROTACs (PROteolysis-TArgeting Chimeras) are small heterobifunctional molecules that induce the degradation of a protein of interest (POI) through the UPS by combining an E3 ubiquitin ligase binding part and a POI binding part in the same molecule connected by a short linker (Sun *et al*, 2019; Sakamoto *et al*, 2001). Ubiquitination is induced when a PROTAC brings together the E3 ligase and the POI, leading to the POI’s ubiquitination and degradation by the proteasome. Although there are more than 600 E3 ligases in humans, only a handful of E3 ligases have been successfully leveraged by PROTACs to induce protein degradation. In some cases, PROTACs fail to induce degradation even when they are found to bind to both E3 and POI (Bondeson *et al*, 2018a; Huang *et al*, 2018). For example, a PROTAC recruiting the E3 ligase VHL was unable to induce the degradation of nonreceptor tyrosine kinases c-Abl and Arg despite formation of a stable ternary complex in cell extracts (Bondeson *et al*, 2018b). It is possible that the PROTACs formed complexes between E3s and POIs but either failed to successfully ubiquitinate the target, or that ubiquitination was successful but led to the placement of the ubiquitin chain in a position that did not allow the proteasome to engage a disordered region in the POI when bound to a ubiquitin receptor on the proteasome. Therefore, keeping this limitation in mind could improve the design of PROTACs and expand the landscape of PROTAC-based therapies.

Taken together, our data demonstrate that the proteasome can effectively degrade substrates with short initiation regions if they are positioned a certain distance from a ubiquitin chain. Moreover, this distance depends on which ubiquitin receptor mediates substrate recognition, likely due to the distance of the ubiquitin receptor to the pore loops which engage the initiation region. Our findings suggest that the ubiquitin chain and disordered region of substrates must be spatially compatible with the architecture of the proteasome to permit their productive engagement and degradation, and may explain why some proteins with a ubiquitin chain and initiation region are not successfully degraded by the proteasome. Understanding the mechanistic constraints that apply to the degradation-targeting signal and its relationship to the architecture of the proteasome sheds light on how the proteasome achieves highly precise substrate selection to maintain cellular functioning. At the same time, two targeting mechanisms with different structural requirements acting in tandem may provide a more robust proteasome function in the face of a highly heterogenous pool of substrates.

## Acknowledgements

These studies were supported by National Institutes of Health grants R01GM124501 to AM, R01GM135264 to AM, and R35 GM145246 to DF, as well as Welch Foundation grant F-1817 to AM.

## Author contributions

CD, BLS, and EC designed and conducted experiments and interpreted data, CD, BLS, EC, JK, HC, and SE constructed unique reagents for the studies, and AM conceived the study, designed experiments, and interpreted data. CD, BLS, and AM wrote the manuscript.

## Materials and Methods

### Molecular biology

#### In Vitro Substrate Proteins

Ub-GFP-tail and UBL-GFP-tail fusion constructs were described previously (Martinez-Fonts & Matouschek, 2016; Martinez-Fonts *et al*, 2020b). Ubiquitin containing constructs, from N- to C-terminus, consist of the coding sequence for *S. cerevisiae* ubiquitin (Ub) with a G76V mutation, a GSGGSG linker, *A. victoria* circular permutant of superfolder GFP (CP8), a G linker, the first 15 or 35 amino acids from *S. cerevisiae* cytochrome *b*_2_ with lysine residues replaced by arginine, and a 6xHis tag. Ubiquitin-like domain (UBL) containing constructs were made by replacing the Ub coding region of the Ub-GFP-tail constructs with the first 80 amino acids of *S. cerevisiae* Rad23. UBL constructs, from N- to C-terminus, consist of UBL derived from Rad23, a GSGGSG linker, *A. victoria* circular permutant of superfolder GFP (CP8), a G linker, the first 15 or 95 amino acids from *S. cerevisiae* cytochrome *b*_2_ with lysine residues replaced by arginine, and a 6xHis tag. Ub-GFP-α-helix-tail and UBL-GFP-α-helix-tail fusion constructs were made by inserting a TS linker, an ER/K α-helix motif of specified amino acid length, followed by a GALM linker between the GFP domain and the tail in the Ub-GFP-tail and UBL-GFP-tail constructs, respectively. Ub-GFP-α-helix(30) was made by removing the tail from Ub-GFP-α-helix(30)-15 and leaving the C-terminal 6xHis tag. UBL-GFP-α-helix(56) was made by removing the tail and 6xHis tag from UBL-GFP-α-helix(56)-15 and moving the 6xHis tag to the N-terminus. Barstar-GFP-α-helix(56)-15 was made by replacing the UBL domain of UBL-GFP-α-helix(56)-15 with the coding region from *E. coli* Barstar protein. All α-helix constructs lacked the C-terminal E244 in GFP to reduce electrostatic interactions between GFP and the Glutamic acid/Glu/E rich α-helix motif. The α-helices were derived from *Mus musculus* myosin VI (Sivaramakrishnan *et al*, 2008). Amino acid sequences of tails and α-helices used in this study are indicated in Appendix Table S3 and Appendix Table S4, respectively. Plasmids used in this study are shown in Appendix Table S2.

### Protein Expression and Purification

#### In Vitro Substrate Proteins

His-tagged GFP containing protein variants were purified as previously described (Martinez-Fonts & Matouschek, 2016), with modifications. Substrate constructs were purified via immobilized metal affinity chromatography through a C-terminal 6xHis-tag by gravity purification. Proteins were expressed in *E. coli* Rosetta (DE3)pLysS (Novagen) cells from pET3a-derived plasmids. A starter culture was inoculated in 25 mL of 2xYT media with 100 ug/mL ampicillin and 34 ug/mL chloramphenicol and grown to OD_600_ of ∼ 0.5 at 37℃ then stored at 4℃ overnight. The starter culture was diluted 1:100 and grown in 2 L of 2xYT media with 100 ug/mL ampicillin and 34 ug/mL chloramphenicol to OD_600_ of ∼ 0.5 at 37℃. Culture was then cooled to 16℃ and induced with 0.4 mM dioxane-free isopropyl B-D-1-thiogalactopyranoside (IPTG) (Calbiochem) overnight at 16℃. Cells were harvested by centrifugation at 5,000xg for 10 min, resuspended in ∼30 mL of NPI-10 (10 mM imidazole, 50 mM phosphate buffer pH 8, 300 mM NaCl), and stored at −80C. For purification, cold NPI-10 buffer was added up to 50 mL. Cells were supplemented with 1X Protease Inhibitor Cocktail Set V EDTA-Free (Calbiochem), 10 mM MgCl_2_, and DNase I (MP Biomedical) and thawed in a room temperature water bath. Cells were then lysed between 10,000 and 15,000 psi in a high-pressure homogenizer (EmulsiFlex-C3 homogenizer, Avestin) for 10 minutes at 4℃. Lysate was cleared by centrifugation at 38,400xg for 20 min at 4℃. Clarified lysate was then filtered through a 0.45 µm syringe filter and mixed with 2 mL of washed TALON® Metal Affinity Resin beads (Takara). The mixture was allowed to bind under nutation at 4℃ for 1 hr. The mixture was poured into a PD-10 column and allowed to settle by gravity. The column was washed once with 5 CV (column volume) of NPI-10 and once with 10 CV of NPI-35 (35 mM imidazole, 50 mM phosphate buffer pH 8, 300 mM NaCl). Protein was eluted three times with 1 CV of NPI-250 (250 mM imidazole, 50 mM phosphate buffer pH 8, 300 mM NaCl). The eluate was concentrated between 1 and 1.5 mL and buffer exchanged in an Amicon Ultra Centrifugation filter with a 10 kDa cutoff into 50 mM TrisHCl of pH at least 1 unit above pI of the protein, 50 mM NaCl, and 0.5 mM DTT. Protein concentration was determined by Pierce Assay. Glycerol was added to 5%, protein aliquoted, flash frozen using liquid nitrogen, and stored at −80℃. Purity was assessed by SDS-PAGE analysis. Proteins with 95 amino acid length initiation regions were purified as described above expect with the addition of 1 mM PMSF to thawed cells.

#### Ubiquitin and Ubiquitin Variants

His6-HRV3C-Ub(K48R) was purified as described previously (Martinez-Fonts & Matouschek, 2016) by immobilized metal affinity chromatography through an N-terminal 6xHis-tag using a HisTrap FF Crude 5 mL column (GE) on an FPLC system (AKTAPurifier). Protein was expressed and lysed as described above for substrate constructs. Lysate was cleared by centrifugation at 38,400xg for 20 min at 4℃ two times. Clarified lysate was then filtered through a 0.45 µm syringe filter and applied to a 5 mL HisTrap FF Crude column on an FPLC system at a rate of 3 mL/min. The column was washed with 5 CV of NPI-10, washed with 10 CV of NPI-20 (20 mM imidazole, 50 mM phosphate buffer pH 8, 300 mM NaCl), and protein eluted with 10 CV of NPI-250 in 2 mL fractions. Fractions containing protein were pooled, concentrated between 1 and 2 mL, and buffer exchanged in an Amicon Ultra Centrifugation filter with a 3 kDa cutoff into 50 mM TrisHCl pH 7.6 and 50 mM NaCl. Protein concentration was determined by Pierce Assay. The protein was aliquoted, flash frozen using liquid nitrogen, and stored at - 80℃. Purity was assessed by SDS-PAGE analysis.

Ubiquitin (*S. cerevisiae*) was purified using published methods (Pickart & Raasi, 2005) by cation exchange chromatography using a Resource S 6 mL column (GE) on an FPLC system (AKTAPurifier). Ubiquitin was expressed as described above except that the 2 L culture was induced with 0.4 mM dioxane-free isopropyl B-D-1-thiogalactopyranoside (IPTG) (Calbiochem) for four hrs at 37℃. Cells were harvested by centrifugation at 5,000xg for 10 min, resuspended in ∼30 mL of 50 mM TrisHCl pH 7.6 and stored at −80℃. For purification, cold 50 mM TrisHCl pH 7.6 was added up to 50 mL. Cells were supplemented with 1X Protease Inhibitor Cocktail Set V EDTA-Free (Calbiochem), 10 mM MgCl_2_, and DNase I (MP Biomedical) and thawed in a room temperature water bath. Cells were then lysed between 10,000 and 15,000 psi in a high-pressure homogenizer (EmulsiFlex-C3 homogenizer, Avestin) for 10 minutes. Lysate was cleared by centrifugation at 38,400xg for 20 min at 4℃. Perchloric acid was added to clarified lysate (0.5% (v/v)) and precipitated proteins removed by centrifugation at 38,400xg for 20 min at 4℃. Supernatant was dialyzed against 2 L of 50 mM ammonium acetate pH 4.5 at 4℃ for four hours in SnakeSkin® dialysis tubing with a 3.5 kDa cutoff (ThermoScientific). Dialysis was repeated in 2 L of fresh buffer overnight at 4℃. The dialysate was filtered through a 0.45 µm syringe filter then applied to a Resource S 6 mL column (GE). Column was washed with 5 CV of 50 mM ammonium acetate pH 4.5 and ubiquitin eluted by applying a linear gradient of 0 to 500 mM NaCl in 50 mM ammonium acetate pH 4.5 over 20 CV in 2 mL fractions. Fractions containing ubiquitin were pooled, concentrated between 1 and 2 mL, and buffer exchanged in an Amicon Ultra Centrifugation filter with a 3 kDa cutoff into 50 mM TrisHCl pH 7.6 and 50 mM NaCl. Concentration was determined by Pierce Assay, protein aliquoted, flash frozen using liquid nitrogen, and stored at −80℃. Purity was assessed by SDS-PAGE analysis.

#### Ubiquitination enzymes

Ube1 (*Mus musculus*) was purified as described previously (Carvalho *et al*, 2012) by immobilized metal affinity chromatography through an N-terminal 6xHis-tag by gravity purification with some modifications described. Ube1 was expressed in *E. coli* Rosetta (DE3)pLysS (Novagen) cells from a pET28-derived plasmid. A starter culture was inoculated in 50 mL of LB media with 25 ug/mL kanamycin and 34 ug/mL chloramphenicol and grown to OD_600_ of ∼ 0.6 at 37℃ then stored at 4℃ overnight. The starter culture was diluted 1:200 and grown in 1 L of LB media with 25 ug/mL kanamycin and 34 ug/mL chloramphenicol to OD_600_ of ∼ 0.6 at 37℃. Culture was then cooled to 16℃ and induced with 0.5 mM dioxane-free isopropyl B-D-1-thiogalactopyranoside (IPTG) (Calbiochem) overnight at 16℃. Cells were harvested by centrifugation at 5,000xg for 10 min, resuspended in ∼30 mL of NPI-10 and stored at −80℃. For purification, cold NPI-10 was added to 50 mL. Cells were supplemented with 0.1% TritonX-100, 1 mM DTT, 1X Protease Inhibitor Cocktail Set V EDTA-Free (Calbiochem), 10 mM MgCl_2_, and DNase I (MP Biomedical) and thawed in a room temperature water bath. Cells were then lysed between 10,000 and 15,000 psi in a high-pressure homogenizer (EmulsiFlex-C3 homogenizer, Avestin) for 10 minutes at 4℃. Lysate was cleared by centrifugation at 38,400xg for 20 min at℃. Clarified lysate was then filtered through a 0.45 µm syringe filter and mixed with 1.5 mL of washed Ni-NTA Agarose beads (Qiagen). The mixture was allowed to bind under nutation at 4℃ for 2 hr. The mixture was poured into a PD-10 column and allowed to settle by gravity. The column was washed three times with 10 CV of NPI-10, once with 10 CV of NPI-20, and eluted three times with 1 CV of NPI-250. The eluate was concentrated to ∼1.0 mL and buffer exchanged in an Amicon Ultra Centrifugation filter with a 30 kDa cutoff into 10 mM TrisHCl pH 8, 1 mM EDTA, and 1 mM DTT. Protein concentration was determined by Pierce Assay and assumed 40% pure as described in published protocols (Carvalho *et al*, 2012). Protein was aliquoted, flash frozen using liquid nitrogen, and stored at −80℃.

E2-25K (*Homo sapiens*) was purified by affinity chromatography as a GST-fusion protein by gravity purification and GST-tag removed by GST-HRV3C protease as described by the published protocol (Cannon *et al*, 2015) with some modifications. E2-25K was expressed in *E. coli* Rosetta (DE3)pLysS (Novagen) cells from a pGEX-6P-1-derived plasmid. A starter culture was inoculated in 25 mL of 2xYT media with 100 ug/mL ampicillin and 34 ug/mL chloramphenicol and grown to OD_600_ of ∼ 0.6 at 37℃ then stored at 4℃ overnight. The starter culture was diluted 1:100 and grown in 2 L of 2xYT media with 100 ug/mL ampicillin and 34 ug/mL chloramphenicol to OD_600_ of ∼ 0.6 at 37℃. Culture was then cooled to 16℃ and induced with 0.4 mM dioxane-free isopropyl B-D-1-thiogalactopyranoside (IPTG) (Calbiochem) overnight at 16℃. Cells were harvested by centrifugation at 5,000xg for 10 min, resuspended in ∼30 mL of 1X PBS and stored at −80℃. For purification, cold 1X PBS was added to 50 mL. Cells were supplemented with 1% TritonX-100, 1 mM DTT, 1X Protease Inhibitor Cocktail Set V EDTA-Free (Calbiochem), 10 mM MgCl_2_, and DNase I (MP Biomedical) and thawed in a room temperature water bath. Cells were then lysed between 10,000 and 15,000 psi in a high-pressure homogenizer (EmulsiFlex-C3 homogenizer, Avestin) for 10 minutes at 4℃. Lysate was cleared by centrifugation at 38,400xg for 20 min at 4℃. Clarified lysate was then filtered through a 0.45 µm syringe filter and mixed with 2.5 mL of washed Glutathione Sepharose 4B beads (GE). The mixture was allowed to bind under nutation at 4℃ for 2 hr. The mixture was poured into a PD-10 column and allowed to settle by gravity. The column was washed three times with 10 CV of 1X PBS supplemented with 1 mM DTT then washed once with 10 CV of GST HRV3C cleavage buffer (150 mM NaCl, 50 mM TrisHCl pH 7.4, 1 mM EDTA, 1 mM DTT). The column was filled with GST HRV3C cleavage buffer and incubated with GST-HRV3C Protease overnight at 4℃. E2-25K was eluted twice with 3 CV of GST HRV3C cleavage buffer. Eluate was concentrated between 1 and 1.8 mL and buffer exchanged in an Amicon Ultra Centrifugation filter with a 10 kDa cutoff into 50 mM TrisHCl pH 7.6, 50 mM NaCl, and 0.5 mM DTT. Protein concentration was determined by Pierce Assay. Glycerol was added to 5%, protein aliquoted, flash frozen using liquid nitrogen, and stored at −80℃. Purity was assessed by SDS-PAGE analysis.

#### Affinity Tag Protease

His6-HRV3C Protease was purified by immobilized metal affinity chromatography through an N-terminal 6xHis-tag by gravity purification. Protein was purified as described above for His-tagged substrate constructs.

### Ubiquitin Chain Synthesis

Ub_4_(K48) and Ub_8_(K48) ubiquitin chains were generated using enzymes that are part of the natural synthesis machinery and purified following the published protocols (Martinez-Fonts & Matouschek, 2016; Cannon *et al*, 2015) as described below. Chains were synthesized from a mixture of wildtype ubiquitin and an N-terminally 6xHis-tagged ubiquitin in which lysine residue 48 was mutated to arginine so chain synthesis terminated with the His-tagged ubiquitin molecule. Polyubiquitin chains of different lengths were then isolated in three steps: an initial enrichment using the N-terminal 6xHis-tag (elution with His6-HRV3C protease) was followed by purification on a cation exchange column (Resource S) using a linear salt gradient and finally size exclusion chromatography (Superdex 75).

K48-linked ubiquitin chains were generated using *H. sapiens* ubiquitin-conjugating enzyme E2-25K together with *the M. musculus* ubiquitin activating enzyme (E1) Ube1 acting on a mixture of wildtype *S. cerevisiae* ubiquitin and a mutant of S. cerevisiae ubiquitin carrying the mutation K48R and an N-terminal 6xHis-tag attached through a linker containing a HRV3C protease cleavage site (His-HRV3C-Ub(K48R)). Chains of four ubiquitin moieties in length were synthesized by incubating 7.5 mg mL^−1^ wildtype ubiquitin and 7.5 mg mL^−1^ His-HRV3C-Ub(K48R), in one-fifth volume of PBDM8 buffer (250 mM Tris-HCl pH 8.0, 25 mM MgCl_2_, 50 mM creatine phosphate, 3 units mL^−1^ inorganic pyrophosphatase, 3 units mL^−1^ creatine phosphokinase), 2.5 mM ATP, and 0.5 mM DTT, with 30 μM E2-25K, and 0.4 μM E1 at 37℃ overnight. The reactions were quenched with 5 mM DTT and aggregates were removed by centrifugation at 15,000 x g for 5 min at 4℃. The supernatants were then diluted with an equal volume of NPI-10, 1 mL of washed Ni-NTA beads (Qiagen catalog no. 30210) were added for each 50 mg of total ubiquitin in the reaction, and the mixtures were allowed to bind under nutation at 4 °C for 1 h. The mixture was then poured into an empty PD-10 column and allowed to settle by gravity. The column was washed twice with 10 CV of NPI-10, followed by 10 CV of HRV3C cleavage buffer (50 mM Tris HCl pH 7.4, 150 mM NaCl, 1 mM DTT). His6-HRV3C protease in HRV3C cleavage buffer was then added to the column, the column capped and nutated overnight at 4℃. Finally, the chains were eluted with 1 CV HRV3C cleavage buffer. For the next purification step by cation exchange chromatography, the eluate was acidified by the addition of 0.03 volumes of 2 N acetic acid to a pH of 4 and loaded on to 6 mL Resource S (GE catalog no. 17-1180-01) in a Tricorn column equilibrated with 50 mM ammonium acetate pH 4.5. The column was washed with 2 CV of 50 mM ammonium acetate pH 4.5 and the chains were eluted in the same buffer with a NaCl gradient as follows: 1 CV of 0 to 200 mM NaCl, 25 CV of 200 to 450 mM NaCl, and 1 CV of 450 to 1000 mM NaCl in 50 mM ammonium acetate pH 4.5, taking 2 mL fractions. The peak containing the chains of the desired length was concentrated to 0.5 mL using an Amicon Ultra Centrifugation filter with a 3 kDa molecular weight cutoff. The chains were purified further on a Superdex Hi-Load 75-pg (GE catalog no. 28-9893-33) column at a rate of 0.25 mL min^−1^ in ubiquitin size exclusion buffer (150 mM NaCl, 50 mM Tris-HCl pH 7.6, 0.5 mM EDTA), collecting 2 mL fractions. Fractions containing chains of the desired length were concentrated and exchanged into 50 mM Tris-HCl pH 7.6 using an Amicon Ultra Centrifugation filter with a 3 kDa molecular weight cutoff. Chain concentration was determined by Pierce Assay. Protein was aliquoted, flash frozen using liquid nitrogen, and stored at −80℃. Purity was assessed by SDS-PAGE analysis.

### Ubiquitinated Substrate Synthesis

Ubiquitinated substrates were generated following the published protocol (Martinez-Fonts & Matouschek, 2016). Polyubiquitin chains were attached to the ubiquitin domain in the target protein using the same enzymes used to create the polyubiquitin chains. After incubation with enzymes and free ubiquitin chains, the reaction was quenched with 5 mM DTT and aggregates were removed by centrifugation at 15,000 x g for 5 min at 4 °C. Base protein was separated from unreacted ubiquitin chains and enzymes by nickel affinity chromatography using the C-terminal His-tag on the base protein. The reaction was mixed with 1.5 mL of washed Ni-NTA Agarose beads (Qiagen). The mixture was allowed to bind under nutation at 4℃ for 1 hr. The mixture was poured into a PD-10 column and allowed to settle by gravity. The column was washed once with 5 CV of NPI-10 and once with 10 CV of NPI-20. Protein was eluted three times with 1 CV of NPI-250. Finally, modified and unmodified base protein were separated by size exclusion chromatography. The eluate from the nickel column was concentrated to 0.5 mL using an Amicon Ultra Centrifugation filter with a 30 kDa molecular weight cutoff and applied to a Superdex Hi-Load 200-pg (GE catalog no. 28-9893-35) column at a flow rate of 0.25 mL min^−1^ in ubiquitin size exclusion buffer supplemented with 1 mM DTT, collecting 2 mL fractions. Fractions containing ubiquitinated substrate were pooled, concentrated, and buffer exchanged into 50 mM Tris-HCl pH 7.6 and 0.5 mM DTT using an Amicon Ultra Centrifugation filter with a 30 kDa molecular weight cutoff. Protein concentration was determined by Pierce Assay. Glycerol was added to 5%, protein aliquoted, flash frozen using liquid nitrogen, and stored at −80℃. Purity was assessed by SDS-PAGE analysis.

### Proteasome Purification

Proteasome was purified from *S. cerevisiae* following published protocols (Sone *et al*, 2004) with modifications described below. The Core Particle (CP) and Regulatory Particle (RP) of the proteasome were purified separately using 3xFLAG tags fused to Pre1 and Rpn11, respectively. The CP was purified from YYS37 (Sone *et al*, 2004) and the regulatory particles were purified from strains indicated in Appendix Table S1. Strains created for this study were constructed using standard methods described (Shi *et al*, 2016a) and are isogenic SUB61 (*MAT α lys2-801 leu2-3,2-112 ura3-52 his3-Δ200 trp1-1(am)* (Finley *et al*, 1987).

Starter yeast cultures were inoculated into 5 mL of YPD with 2% glucose and grown for 24 hrs at 30℃. Culture was diluted 1:100 into 2 L of YPD with 2% glucose and grown to OD_600_ ∼ 2 at 30℃. Cells were centrifuged at 5,000xg for 10 min, washed with cold water, and centrifuged again at 5,000xg for 10 min. Cells were then washed in 1X Buffer A (50 mM TrisHCl pH 7.6, 5% glycerol), centrifuged at 5,000xg for 10 min, and stored at −80℃. For purification, cells were resuspended in ARS Wash Buffer (50 mM TrisHCl pH 7.6, 5% glycerol, 1 mM ATP, 10 mM MgCl_2_, 1 mM DTT, 20 mM creatine phosphate, 0.02 mg/mL creatine phosphokinase). Cells were lysed between 25,000 and 30,000 psi in a high-pressure homogenizer (EmulsiFlex-C3 homogenizer, Avestin) for 10 minutes at 4℃. Lysates were clarified by centrifuging at 30,000xg for 30 min at 4℃. Clarified lysate was filtered with a 0.45 µm filter syringe then supplemented with 5 mM ATP, 10 mM creatine phosphate, and 0.01 mg/mL creatine phosphokinase. Supplemented lysate was then mixed with 1 mL of pre-washed anti-FLAG M2 agarose beads (Sigma) and allowed to bind under nutation at 4℃ for 2 hrs. The mixture was then collected in a PD-10 column and allowed to settle by gravity. The column was washed twice with 15 CV of ATP Wash Buffer (50 mM TrisHCl pH 7.6, 5% glycerol, 2 mM ATP, 5 mM MgCl_2_, 1 mM DTT) then once with ATP500 buffer (50 mM TrisHCl pH 7.6, 5% glycerol, 2 mM ATP, 5 mM MgCl_2_, 1 mM DTT, 500 mM NaCl). The column was filled with ATP500 buffer, capped, and nutated for 1 hr at 4℃. Mixture was allowed to settle by gravity, washed twice with 15 CV of ATP500 buffer, washed twice with 15 CV of ATP Wash Buffer, and excess buffer was removed by centrifuging at 500xg for 5 sec. Column was incubated with 0.75 mL of Elution Buffer (0.15 mg/mL 3xFLAG peptide in ATP Wash Buffer) for 20 min at room temperature and proteasome eluted by centrifuging at 500xg for 5 sec. A second elution was performed using 0.5 mL Elution Buffer, and the two elutions were pooled. Concentration was determined by Pierce Assay. The concentration of the Elution Buffer was subtracted from the concentration of the pooled eluates to determine the proteasome concentration. Eluate was aliquoted, flash frozen in liquid nitrogen, and stored at −80℃. Proteasome composition and purity was assessed by SDS-PAGE analysis.

### Kinetic Plate Reader Degradation Assays

Single turnover degradation assays were performed as previously described (Martinez-Fonts *et al*, 2020b) with modifications. In short, 25 nM reconstituted proteasome was presented to 5 nM purified substrates in the presence of an ATP Regeneration System (ARS) at 30℃ in 384-well plates (flat bottom, low flange, non-binding, black, Corning).

Proteasomes were first reconstituted in a 1:2 ratio at twice that of the final concentration by incubating 50 nM CP and 100 nM RP in presence of 2X ARS (2 mM ATP, 20 mM creatine phosphate, 0.2 mg/mL creatine phosphokinase), 1X Degradation Buffer (50 mM TrisHCl pH 7.6, mM MgCl_2_, 1% glycerol), and 8 mM DTT at 30℃ for 30 min. Substrates were diluted to twice that of the final concentration (i.e., 10 nM) using protein buffer (4 mg/mL BSA, 1X Degradation Buffer, 100 mM NaCl). Reactions were initiated by mixing 20 µL of proteasome mixture with 20 µL of substrate mixture. Thus, final reactions contained 25 nM proteasome and 5 nM substrate in 1X ARS (1 mM ATP, 10 mM creatine phosphate, 0.1 mg/mL creatine phosphokinase), 4 mM DTT, 1X Degradation Buffer, 2 mg/mL BSA, and 50 mM NaCl.

Substrate fluorescence (488 nm excitation, 520 nm emission) was measured every 60 seconds for 90 minutes in a plate reader (Infinite M1000 PRO, Tecan). Background (average fluorescence of well containing everything except substrate) was subtracted from proteasome containing reactions and fluorescence plotted as a function of time. Likewise, background (average fluorescence of well containing everything except substrate and proteasome) was subtracted from reactions lacking proteasome and fluorescence plotted as a function of time. The decay curves were fitted to the equation describing a single exponential decay (y = Ae^-kt^ + C) using the software Prism (version 7). Initial degradation rate was determined by multiplying the amplitude of degradation (% fluorescence) by the rate constant (min^-1^).

### Yeast Model Substrate Design and Expression

Yeast YFP substrate variants were built off a central YFP (sYFP2) domain and constructed with either an N-end rule degron, or an N-terminal UBL domain derived from the first 80 amino acids of *S. cerevisiae* Rad23. N-end rule substrates consisted of a ubiquitin domain followed by a destabilizing Arg residue followed by a 19 amino acid linker derived from *E. coli* lacI, which contains two Lys residues (Rekk). However, amino acids 4-13 were replaced with a serine-rich linker derived from herpes virus 1 ICP4 to prevent the N-terminal degron from being a disordered region that could initiate degradation. To correct for cell-to-cell variation in transcriptional, translational, or cell size variation, RFP dsRed-Express2 was expressed upstream of YFP separated by a P2A ribosomal skipping site. The ratio of YFP over RFP for individual cells was a measure of the steady state concentration of YFP variants.

The N-end rule degron and UBL yeast substrates with Su9 and SP25 tails were individually expressed from CEN plasmids using a constitutive *TPI1* promoter. The Su9 tail represents 51 amino acids from subunit 9 of the F_o_ component of the *Neurospora crassa* ATP synthase and has been previously characterized as an efficient initiation region (Yu *et al*, 2016b). The SP25 tail consists of 5 repeats of peptide region 2 in influenza A virus M2 protein used to produce antisera and has been previously characterized to be a poor initiation region (Yu *et al*, 2016b). Plasmids were then transformed into WT, *ubr1*Δ, and *pep4*Δ yeast strains. Plasmids used in this study are shown in Appendix Table S2. The deletion strains were obtained from the yeast gene deletion collection (Giaever *et al*).

Ubiquitinated and UBL substrates were depleted when attaching a tail that is recognized by the proteasome (Su9, Fig EV4a) (Yu *et al*, 2016a), but accumulated when attaching a tail that is not recognized (SP25, Fig EV4a). Deleting the primary ubiquitin ligase that targets N-end rule substrates (*ubr1*Δ) to the proteasome caused buildup of substrate with the Su9 tail, while not affecting degradation of UBL substrate, indicating N-end rule substrates require ubiquitination for degradation. Deleting the gene encoding proteinase A (*pep4*Δ), which functions in the final step of autophagy, did not affect substrate abundance (Fig EV4b) suggesting substrates are not degraded through the autophagy-lysosome pathway. Taken together, the YFP model substrate platform is an effective way to measure degradation by the proteasome in yeast.

Varying amino acid lengths of ER/K α-helix motifs were inserted between YFP and tail of N-end rule degron and UBL substrates to generate YFP spacing substates. The tail was the same as the 18 amino acid long tail attached to GFP substrates used in the *in vitro* studies, but is 19 amino acids in length due to insertion of a glycine/Gly/G residue at position three. Tails retained a C-terminal 6xHis tag to maintain design consistency between *in vitro* and *in vivo* substrates.

Substrates containing α-helices were integrated into the ORF of HO1 in *S. cerevisiae* and expressed from a constitutive *TPI1* promoter. Correct integration was confirmed by selection with nourseothricin and sanger sequencing. Yeast strains used in this study are shown in Appendix Table S5.

### Flow Cytometry

Cells were grown in synthetic complete medium at 25°C to mid-log phase after 7-8 doublings and harvested for direct fluorescent measurement of RFP and YFP channels by an LSR Fortessa (BD biosciences) flow cytometer. Time courses of YFP fluorescence were measured to determine *in vivo* degradation rates by inhibition of protein synthesis through the addition of 125 µM cycloheximide and YFP fluorescence measurements were taken for 120 minute time courses. Degradation rates were calculated in the same way as in the *in vitro* experiments. The data were analyzed by FlowJo (FlowJo version 10.2). Cells were gated first along the FSC-A vs. SSC-A axes to eliminate dead cells. They were then gated against SSC-H vs. SSC-W and FSC-H vs. FSC-W axes to eliminate doublets. Finally, cells were gated to only include RFP-positive cells. YFP over RFP ratios of at least 10,000 cells were calculated and we report the median of those ratios for each experiment.

### Data Availability

This study includes no data deposited in external repositories.

## Expanded View Figures

**Figure EV1.**
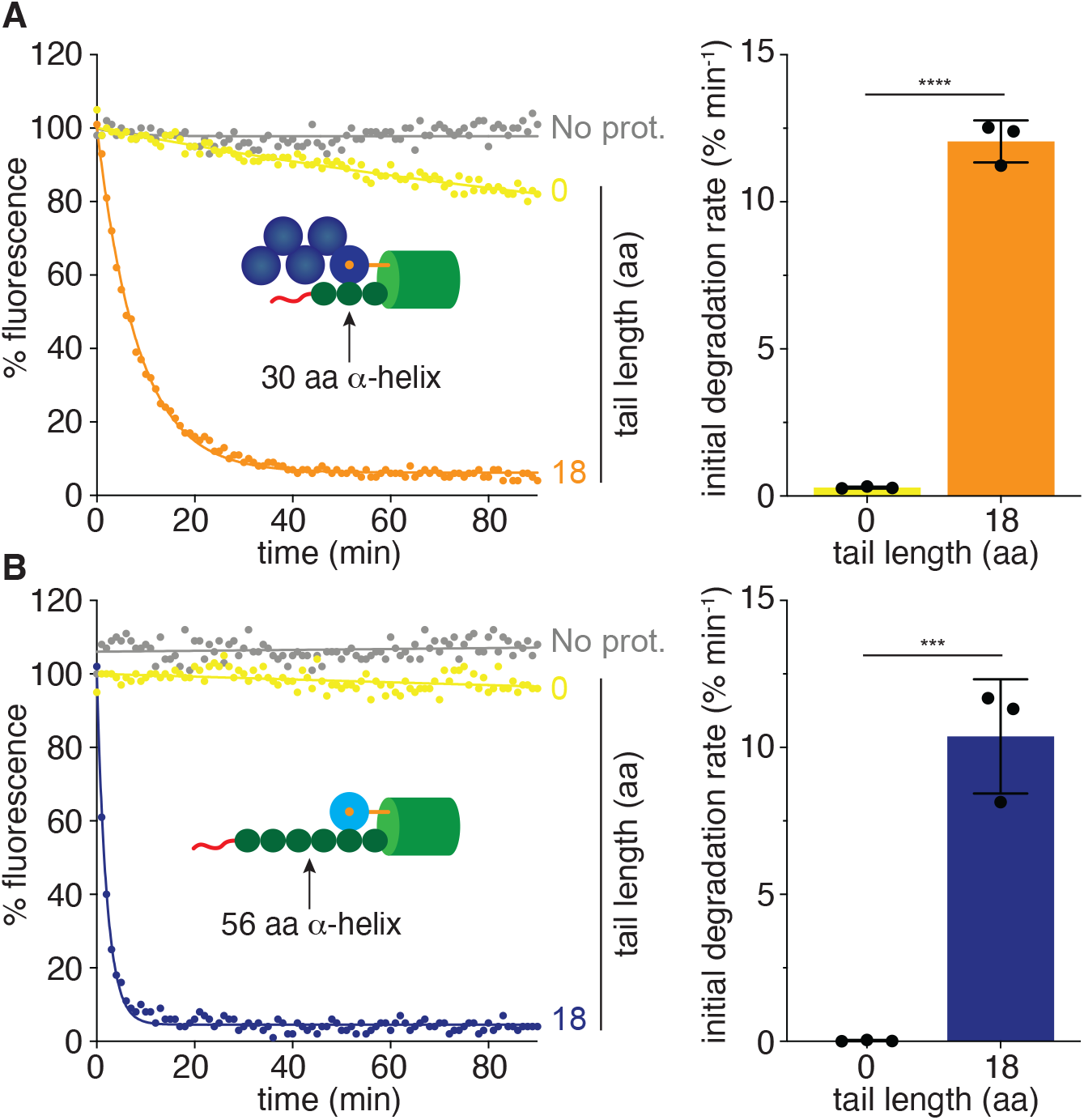
The α-helices do not function as effective initiation regions. A (Left) *In vitro* degradation kinetics of Ub_5_(K48)-tagged GFP substrate with the 30 amino acid (aa) long α-helix linker and either a 15 (orange) or 0 aa length tail (yellow) under single turnover conditions (5 nM substrate, 25 nM wildtype (WT) proteasome). Substrate fluorescence is plotted as a percentage of the initial fluorescence as a function of time in minutes. No proteasome (no prot) control included. (Right) Initial degradation rates of substrates with the indicated tail calculated by fitting kinetics to single exponential decay. B (Left) *In vitro* degradation kinetics of UBL-tagged GFP substrate with the 56 amino acid (aa) long α-helix linker and either a 15 tail (blue) or 0 aa length tail (yellow) under single turnover conditions (5 nM substrate, 25 nM WT proteasome). Substrate fluorescence is plotted as a percentage of the initial fluorescence as a function of time in minutes. (Right) Initial degradation rates of substrates with the indicated tail calculated by fitting kinetics to single exponential decay. Data information: In (A,B), experiments were performed in triplicate, one representative kinetic experiment shown. In (A,B) rate data represent the means from three experiments and error bars represent the standard deviations (SD). **** P < 0.0001 and *** P < 0.001 (two-tailed unpaired t test).

**Figure EV2.**
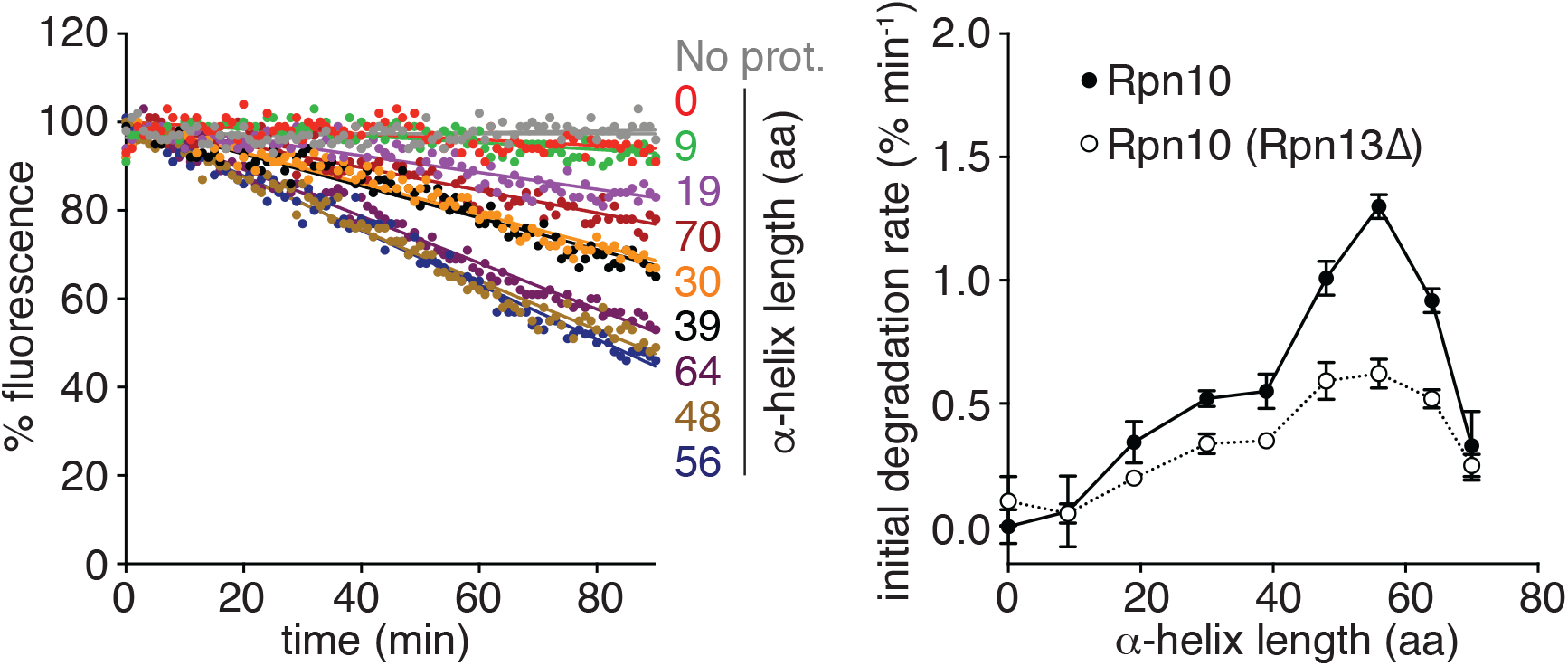
Degradation of UBL-tagged spacing substrates on Rpn10 only proteasome is due to residual binding to mutated Rpn13. (Left) *In vitro* degradation kinetics of UBL spacing substrates under single turnover conditions (5 nM substrate, 25 nM proteasome) by Rpn10(*Rpn13*Δ) proteasome. Graph plots substrate fluorescence as a percentage of the initial fluorescence as a function of time in minutes. The amino acid (aa) length of the α-helix is shown to the right of the corresponding curve. (Right) Initial degradation rates of UBL spacing substrates by the indicated proteasome were calculated by fitting kinetics to single exponential decay or linear equation. Graph plots the rate as a function of aa length of the α-helix. Data information: experiments were performed in triplicate, one representative kinetic experiment shown. Rate data are plotted as a mean calculated from at least three experiments +/- the SD. Proteasome types are described in Appendix Table S1.

**Figure EV3.**
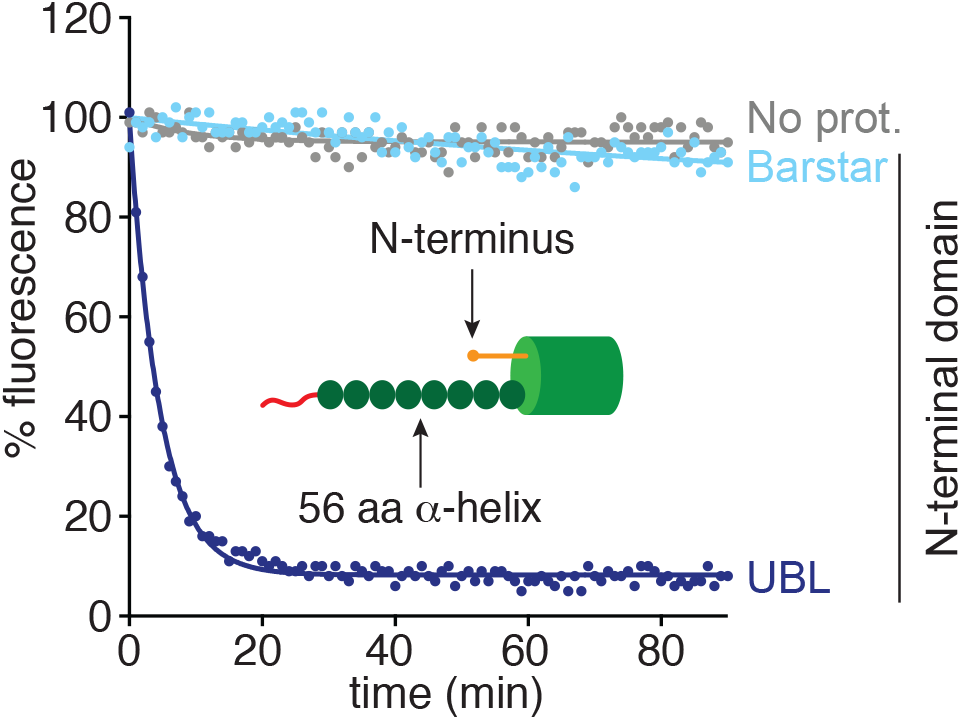
Degradation of UBL-tagged substrate is due to UBL binding the proteasome. *In vitro* degradation kinetics of model GFP substrates with the 56 amino acid (aa) length α-helix, 18 tail, and the indicated N-terminal domain under single turnover conditions (5 nM substrate, 25 nM wildtype (WT) proteasome). Graph plots substrate fluorescence as a percentage of the initial fluorescence as a function of time in minutes. The N-terminal domain is shown to the right of the corresponding curve. Data information: experiment was performed in triplicate, one representative kinetic experiment shown.

**Figure EV4.**
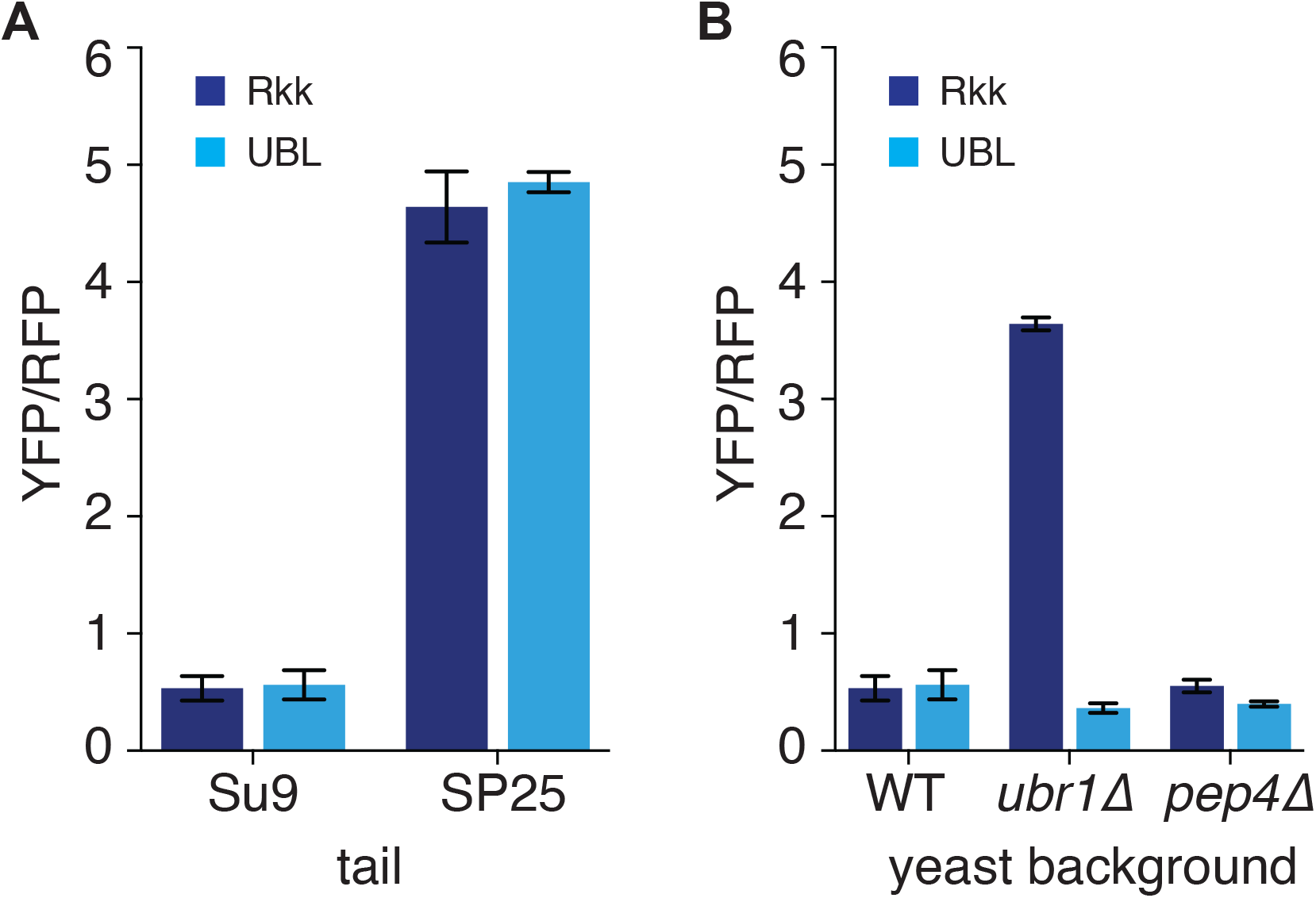
Model YFP proteins are proteasome substrates and degraded in a ubiquitin-dependent and autophagy-independent manner. A Corrected medians of cellular YFP fluorescence (median YFP/RFP values) representing steady-state degradation of yeast cultures expressing YFP substrates targeted to the proteasome through an N-end degron (Rkk, dark blue) or UBL domain (cyan) with Su9 or SP25 tail. Proteasome substrates were expressed in the genetic background *by4741* (WT, wild type) B Corrected medians of cellular YFP fluorescence (median YFP/RFP values) representing steady-state degradation of yeast cultures expressing YFP substrates targeted to the proteasome through an N-end degron (Rkk, dark blue) or UBL domain (cyan) with Su9 tail. Proteasome substrates were expressed in the genetic background *by4741* with no additional mutations (WT, wild type), the open reading frame of UBR1 deleted (*ubr1*Δ) to abolish ubiquitination by the N-end rule pathway, or the open reading frame of PEP4 deleted (*pep4*Δ) to reduce proteolytic activity involved in autophagy. Data information: experiments were performed in triplicate. YFP/RFP ratios are plotted as a mean calculated from at least three experiments +/- the SD. Yeast strains are described in Appendix Table S5.

