## Supplemental Tables for "A strict requirement in proteasome substrates for spacing between ubiquitin tag and degradation initiation elements"

**for**

**[1]Department of Molecular Biosciences, The University of Texas at Austin, Austin, TX**

**[2]Department of Cell Biology, Harvard Medical School, Boston, MA**

**Table of Contents**


Tables……………………………………………………………………………………………… 3

Appendix Table S1…………………………………………………………………………….. 3

Appendix Table S2…………………………………………………………………………….. 4

Appendix Table S3…………………………………………………………………………….. 6

Appendix Table S4…………………………………………………………………………….. 7

Appendix Table S5…………………………………………………………………………….. 9

References………………………………………………………………………………………. 10

**Tables**

Appendix Table S1. Proteasome mutant yeast strains used in this study.

| Strain Name | Genotype | Proteasome Label |
|---|---|---|
| YYS40 * | *MAT α RPN11-FLAG::HIS3* | RP (wildtype) |
| YYS37 * | *MAT α PRE1-FLAG::HIS3* | CP |
| SY1962a | *MAT α RPN11-FLAG::HIS3 rpn1-AKAA-ARR::HGR rpn13-PRU::NATmx* | Rpn10 |
| SY1961a | *MAT α RPN11-FLAG::HIS3 rpn1-AKAA-ARR::HGR rpn10-UIM::KANmx* | Rpn13 |
| SY1572b | *MAT α RPN11-FLAG::HIS3 rpn10-UIM::KANmx rpn13-PRU::NATmx* | Rpn1 |
| SY1960a | *MAT α RPN11-FLAG::HIS3 rpn1-AKAA-ARR::HGR rpn10-UIM::KANmx rpn13-PRU::NATmx UBP6-pADH1-Ub::URAmx* | QM (quadruple mutant) |
| BLS635 | *MAT a RPN11-FLAG::HIS3 rpn1-AKAA-ARR:: rpn13::KANmx4* | Rpn10 (Rpn13Δ) |

*Strains published in ref (Sone *et al*, 2004). All other strains were created for this study and are isogenic to SUB61 (*MAT α lys2-801 leu2-3,2-112 ura3-52 his3- Δ200 trp1-1 (am)* ref (Finley *et al*, 1987)).

The *rpn10-uim* allele carries the mutations L228N, A229, M230N, A231N, and L232N ref (Fu *et al*, 1999). The *rpn13-pru* allele carries the mutations E41K, E42K, L43A, F45A, and S93D ref (Shi *et al*, 2016). The *rpn1-AKAA-ARR* allele carries the T1 site mutations D541A, D548R, and E552R and T2 mutations L430A, D431K, Q434A, Q435A ref (Shi *et al*, 2016). Strain SY1960a carries an additional ubiquitin gene driven by the strong ADH1 promoter and integrated at the UBP6 locus to compensate for poor growth.

Appendix Table S2. Plasmids used in this study

| Plasmid ID Number | Protein | Vector | Function/Use/purpose |
|---|---|---|---|
| pCD#018 | Ub-CP8-35-His6 | pET-3a | Proteasome substrate |
| pCD#023 | Ub-CP8-15-His6 | pET-3a | Proteasome substrate |
| pCD#024 | Ub-CP8-α9-15-His6 | pET-3a | Proteasome substrate |
| pCD#025 | Ub-CP8-α19-15-His6 | pET-3a | Proteasome substrate |
| pCD#041 | Ub-CP8-α30-15-His6 | pET-3a | Proteasome substrate |
| pCD#049 | Ub-CP8-α39-15-His6 | pET-3a | Proteasome substrate |
| pCD#051 | Ub-CP8-α48-15-His6 | pET-3a | Proteasome substrate |
| pCD#053 | Ub-CP8-α56-15-His6 | pET-3a | Proteasome substrate |
| pCD#055 | Ub-CP8-α64-15-His6 | pET-3a | Proteasome substrate |
| pCD#026 | Ub-CP8-α70-15-His6 | pET-3a | Proteasome substrate |
| pCD#073 | Ub-CP8-α30-His6 | pET-3a | Proteasome substrate |
| pCD#071 | UBL(80)-CP8-35-His6 | pET-3a | Proteasome substrate |
| pCD#077 | UBL(80)-CP8-15-His6 | pET-3a | Proteasome substrate |
| pCD#078 | UBL(80)-CP8-α9-15-His6 | pET-3a | Proteasome substrate |
| pCD#079 | UBL(80)-CP8-α19-15-His6 | pET-3a | Proteasome substrate |
| pCD#080 | UBL(80)-CP8-α30-15-His6 | pET-3a | Proteasome substrate |
| pCD#081 | UBL(80)-CP8-α39-15-His6 | pET-3a | Proteasome substrate |
| pCD#082 | UBL(80)-CP8-α48-15-His6 | pET-3a | Proteasome substrate |
| pCD#083 | UBL(80)-CP8-α56-15-His6 | pET-3a | Proteasome substrate |
| pCD#084 | UBL(80)-CP8-α64-15-His6 | pET-3a | Proteasome substrate |
| pCD#085 | UBL(80)-CP8-α70-15-His6 | pET-3a | Proteasome substrate |
| pCD#112 | His6-UBL(80)-CP8-α56 | pET-3a | Proteasome substrate |

| | | | |
|---|---|---|---|
| pCD#037 | Ube1 | pET28 | Ubiquitin chain synthesis |
| pCD#013 | E2-25K | pGEX-6P-1 | Ubiquitin chain synthesis |
| pCD#004 | Ubiquitin (wildtype) | pET-3a | Ubiquitin chain synthesis |
| pCD#030 | His6-HRV3C-Ub(K48R) | pET-3a | Ubiquitin chain synthesis |
| pCD#017 | His6-HRV3C Protease | pETDuet | His-tag cleavage |
| pCD#016 | GST-HRV3C Protease | pGEX-4T-1 | GST-tag cleavage |
| pBLS005 | pTPI1_4A_DsRed_P2A_Ub-Rkk_N10SRR-CΔ20-sYFP-Su9_tADH | YCplac33 | *In vivo* proteasome substrate plasmid |
| pBLS137 | pTPI1_4A_DsRed_P2A_UBL(80)-sYFP-Su9_tADH | YCplac33 | *In vivo* proteasome substrate plasmid |
| pBLS138 | pTPI1_4A_DsRed_P2A_UBL(80)-sYFP-SP25_tADH | YCplac33 | *In vivo* proteasome substrate plasmid |
| pBLS047 | pTPI1_4A_DsRed_P2A_Ub-Rkk_N10SRR-CΔ20-sYFP-SP25_tADH | YCplac33 | *In vivo* proteasome substrate plasmid |

Appendix Table S3. Amino acid sequences of initiation regions (i.e., tails) used in this study.

| Tail Name | Base Substrate | Amino Acid Sequence | Tail Length (amino acids) |
|---|---|---|---|
| 18 | GFP | PRLRYQPLLRSGRISPAE | 18 aa |
| 38 | GFP | PRLRYQPLLRISQNCEAAILRASQTRLNTISGRISPAE | 38 aa |
| 96 | GFP | PRLRYQPLLRISQNCEAAILRASQTRLNTIGAYGSTVPRSQSFEQDSRQRTQSWTALRVGAIPAATSSVAYLNWHNGQIDNEPQLDMNRQRISPAE | 96 aa |
| Su9 | YFP | RMASTRVLASRLASQMAASAKVARPAVRVAQVSKRTIQTGSPLQTRAYSS | 49 aa |
| SP25 | YFP | RSPESMREEYRKEGSPESMREEYRKEGSPESMREEYRKEGSPESMREEYRKEGSPESMREEYRKE | 65 aa |
| 19 | YFP | PRGLRYQPLLRSGRISPAE | 19 aa |

Appendix Table S4. Amino acid sequences of α-helices used in this study.

| **Helix Identification** | **Amino Acid Sequence** |
|---|---|
| α9 | EEEERRRQQ |
| α19 | EEEERRRQQEEEAERLRRI |
| α30 | EEEERRRQQEEEAERLRRIQEEMERERRRR |
| α39 | EEEERRRQQEEEAERLRRIQEEMERERRRREEDEERRRR |
| α48 | EEEERRRQQEEEAERLRRIQEEMERERRRREEDEERRRREEEERRMRL |
| α56 | EEEERRRQQEEEAERLRRIQEEMERERRRREEDEERRRREEEERRMRL EMEARRRQ |
| α64 | EEEERRRQQEEEAERLRRIQEEMERERRRREEDEERRRREEEERRMRL EMEARRRQEEEERRRR |
| α70 | EEEERRRQQEEEAERLRRIQEEMERERRRREEDEERRRREEEERRMRL EMEARRRQEEEERRRREDDERR |

Appendix Table S5. Yeast strains used in this study.

| **Strain Name** | **Genotype** |
|---|---|
| JBK 001 | MATa ho::natMX6_pTPI1_DsRed_P2A_Ub-Rkk_N10SRR-Cd20-sYFP-19-His6_tADH his3Δ1 leu2Δ0 met15Δ0 ura3Δ0 |
| JBK 002 | MATa ho::natMX6_pTPI1_DsRed_P2A_Ub-Rkk_N10SRR-Cd20-sYFP-ahelix09-19-His6_tADH his3Δ1 leu2Δ0 met15Δ0 ura3Δ0 |
| JBK 005 | MATa ho::natMX6_pTPI1_DsRed_P2A_Ub-Rkk_N10SRR-Cd20-sYFP-ahelix19-19-His6_tADH his3Δ1 leu2Δ0 met15Δ0 ura3Δ0 |
| JBK 006 | MATa ho::natMX6_pTPI1_DsRed_P2A_Ub-Rkk_N10SRR-Cd20-sYFP-ahelix30-19-His6_tADH his3Δ1 leu2Δ0 met15Δ0 ura3Δ0 |
| JBK 003 | MATa ho::natMX6_pTPI1_DsRed_P2A_Ub-Rkk_N10SRR-Cd20-sYFP-ahelix39-19-His6_tADH his3Δ1 leu2Δ0 met15Δ0 ura3Δ0 |
| JBK 004 | MATa ho::natMX6_pTPI1_DsRed_P2A_Ub-Rkk_N10SRR-Cd20-sYFP-ahelix48-19-His6_tADH his3Δ1 leu2Δ0 met15Δ0 ura3Δ0 |
| JBK 007 | MATa ho::natMX6_pTPI1_DsRed_P2A_Ub-Rkk_N10SRR-Cd20-sYFP-ahelix56-19-His6_tADH his3Δ1 leu2Δ0 met15Δ0 ura3Δ0 |
| JBK 008 | MATa ho::natMX6_pTPI1_DsRed_P2A_Ub-Rkk_N10SRR-Cd20-sYFP-ahelix64-19-His6_tADH his3Δ1 leu2Δ0 met15Δ0 ura3Δ0 |
| JBK 009 | MATa ho::natMX6_pTPI1_DsRed_P2A_Ub-Rkk_N10SRR-Cd20-sYFP-ahelix70-19-His6_tADH his3Δ1 leu2Δ0 met15Δ0 ura3Δ0 |
| JBK 010 | MATa ho::natMX6_pTPI1_DsRed_P2A_Rad23-sYFP-19-His6_tADH his3Δ1 leu2Δ0 met15Δ0 ura3Δ0 |
| JBK 011 | MATa ho::natMX6_pTPI1_DsRed_P2A_Rad23-sYFP-ahelix09-19-His6_tADH his3Δ1 leu2Δ0 met15Δ0 ura3Δ0 |
| JBK 012 | MATa ho::natMX6_pTPI1_DsRed_P2A_Rad23-sYFP-ahelix19-19-His6_tADH his3Δ1 leu2Δ0 met15Δ0 ura3Δ0 |
| JBK 013 | MATa ho::natMX6_pTPI1_DsRed_P2A_Rad23-sYFP-ahelix30-19-His6_tADH his3Δ1 leu2Δ0 met15Δ0 ura3Δ0 |
| JBK 014 | MATa ho::natMX6_pTPI1_DsRed_P2A_Rad23-sYFP-ahelix39-19-His6_tADH his3Δ1 leu2Δ0 met15Δ0 ura3Δ0 |
| JBK 015 | MATa ho::natMX6_pTPI1_DsRed_P2A_Rad23-sYFP-ahelix48-19-His6_tADH his3Δ1 leu2Δ0 met15Δ0 ura3Δ0 |
| JBK 016 | MATa ho::natMX6_pTPI1_DsRed_P2A_Rad23-sYFP-ahelix56-19-His6_tADH his3Δ1 leu2Δ0 met15Δ0 ura3Δ0 |
| JBK 017 | MATa ho::natMX6_pTPI1_DsRed_P2A_Rad23-sYFP-ahelix64-19-His6_tADH his3Δ1 leu2Δ0 met15Δ0 ura3Δ0 |
| JBK 018 | MATa ho::natMX6_pTPI1_DsRed_P2A_Rad23-sYFP-ahelix70-19-His6_tADH his3Δ1 leu2Δ0 met15Δ0 ura3Δ0 |
| BLS 623 | MATa [pBLS005] his3Δ1 leu2Δ0 met15Δ0 ura3Δ0 |
| BLS 742 | MATa [pBLS005] pep4::kanMX4 his3Δ1 leu2Δ0 met15Δ0 ura3Δ0 |
| BLS 746 | MATa [pBLS005] ubr1::kanMX4 his3Δ1 leu2Δ0 met15Δ0 ura3Δ0 |
| BLS 613 | MATa [pBLS137] his3Δ1 leu2Δ0 met15Δ0 ura3Δ0 |
| BLS 744 | MATa [pBLS137] pep4::kanMX4 his3Δ1 leu2Δ0 met15Δ0 ura3Δ0 |
| BLS 748 | MATa [pBLS137] ubr1::kanMX4 his3Δ1 leu2Δ0 met15Δ0 ura3Δ0 |

| | |
|---|---|
| BLS 611 | MATa [pBLS138] his3Δ1 leu2Δ0 met15Δ0 ura3Δ0 |
| BLS 617-1 | MATa [pBLS047] his3Δ1 leu2Δ0 met15Δ0 ura3Δ0 |

All above strains were created for this study.
